# eIF4E and Ezrin drive a pseudopod-localized migratory translation program in AML

**DOI:** 10.64898/2026.02.21.707190

**Authors:** Biljana Culjkovic-Kraljacic, Leandro Marcelo Martinez, Naiyu Shi, Alyssa Retiz, Samuel Perron, Aksharh Thambipillai, Caleb M. Embree, Winnie Yip, Vicenta Trujillo-Alonso, Emma Lassman, Marysol Chu Carty, Kata Alilovic, Sebastien Carreno, Gail J Roboz, Monica L. Guzman, Katherine L.B. Borden

## Abstract

eIF4E, the mRNA cap-binding protein, is elevated in acute myeloid leukemia (AML) subtypes associated with increased cell movement and leukemic infiltration. We show here that eIF4E is required for AML cell motility, bone marrow engraftment, and disease progression in mouse models, patient specimens and cell line models. While dissecting the underlying mechanisms of eIF4E-dependent motility, we discovered that the motility factor Ezrin, enriched in pseudopods, is required for high-eIF4E AML cell movement. We discovered that eIF4E directly binds Ezrin’s FERM domain and that Ezrin, through this direct interaction with eIF4E, associated with mRNAs encoding its own motility pathway. We developed a method to directly mark active ribosomes *in situ* (VISTA-R) allowing us to observe translation in Ezrin-positive pseudopods. We show that Ezrin anchors actively translating ribosomes in pseudopods, creating a localized translation site we term a T-POD. Depleting Ezrin reduced the number of active ribosomes within T-PODs, without affecting global translation; and Ezrin reduction was sufficient to decrease protein production of factors in the Ezrin-CD44-HA motility pathway corresponding to impaired motility, positioning Ezrin not only as a motility factor but also now as a positive regulator of translation. Furthermore, Ezrin RIP-seq data revealed that Ezrin, through eIF4E, potentially regulates production of many factors involved in invasion including cytoskeletal remodeling and protrusion formation and extracellular matrix remodeling. In all, our studies identify an unanticipated role for Ezrin in a local translation program that sustains AML motility *in vivo* through provisioning T-PODs and revealed an unexpected direct connection between the translation and motility machinery.

## Introduction

Acute myeloid leukemia (AML) carries a poor prognosis, particularly for older adults, who make up the majority of patients. Disease persistence depends on the bone marrow (BM) niche, where leukemic cells adhere to stroma and are sheltered from therapy, and from which they can egress into the circulation to spread to distal BM sites and infiltrate other tissues.^1^ Extramedullary disease, seen in roughly 25–50% of patients, is associated with worse outcomes ^2^. Both retention in and escape from the niche require leukemic cells to move, using amoeboid-like motility driven by pseudopod-like protrusions. The transcriptional and signaling programs that govern AML cell behavior have been studied extensively, but whether post-transcriptional control of gene expression contributes to leukemic cell movement is not well known^3^. Moving cells must also supply new proteins rapidly at their protrusions, raising the possibility that mRNA translation is spatially coordinated with the motility machinery itself. Whether this occurs, and how, has been unclear.

AML cells employ an amoeboid-like strategy reliant on pseudopod-like protrusions for movement through BM stroma whereby the surfaces of these structures are decorated with factors involved in stromal interactions such as the CD44 transmembrane proteins and the glycosaminoglycan hyaluronan (HA) on the stroma^4, 5,6, 7^. Communication between AML cells and the BM matrix is critical for initiating programs to promote migration and subsequent disease progression^1^, where CD44 on the surface of AML cells enables embedding into the HA-rich BM niche; indeed, elevated HA levels correlate with increased adhesion of hematopoietic cells to the niche^7^. The HA-CD44 interaction stimulates signaling events which promote migratory behavior of AML cells and increase production of proteases which in turn degrade cell adhesion junctions thereby generating space through which cells migrate^7^. Additionally, HA binding to CD44 promotes intracellular signaling which drives the mechanics underpinning morphological rearrangements required for cells to invade. CD44 conveys these signals through direct interaction of its intracellular C-terminal tail with the N-terminal ERM domain (Ezrin, Radixin, Moesin) of Ezrin or in some instances with other ERM proteins^8, 9^. Ezrin can simultaneously bind CD44 and through its C-terminal domain the cortical actin. In this way Ezrin directly links CD44 signaling with the machinery driving physical deformations of the membrane required for pseudopod formation and motility of AML and other cancer cells^8, 9^.

Increased leukemic infiltration and extramedullary disease are strongly associated with M4/M5 French American British (FAB) AML subtypes^10^ ^11–14^. Patients with these AML subtypes are characterized by substantially elevated levels and frequent nuclear enrichment of the eukaryotic translation initation factor eIF4E relative to CD34+ BM cells from healthy donors or AML patients with normal eIF4E levels^11–14^. While FAB subtypes are no longer used to classify AML, these data showcase that a considerable fraction of AML patients have elevated eIF4E. Survival analysis of a 609 AML patient cohort (Leucegene.ca) indicated that high-eIF4E AML patients had substantially reduced survival relative to others (p=0.0438)^15^. Given the heterogeneity of AML patients with respect to their various complex karyotypes and mutational landscapes across the analyzed patients, this correlation between eIF4E levels and outcome is strong^15^. Notably, eIF4E did not co-vary in this cohort with other established AML markers such as NPM mutations. Consistent with these findings, another study identified eIF4E elevation in AML cells over healthy volunteers (p<.001) with an enrichment in M5 AML subtypes^10^. Moreover, this study also reported worse outcomes in high-eIF4E AML patients. Additionally, eIF4E also plays roles in HoxA9-dependent AML which are also associated with poorer outcomes than other AMLs^16–18^.

eIF4E influences many aspects of posttranscriptional regulation and translation of specific mRNAs through its engagement of the m^7^G cap. Here, eIF4E uses a molecular cleft to bind these mRNAs directly via their methyl-7-guanosine (m^7^G) “cap” located on the 5’ end of the transcript^19–21^. Highlighting selectivity, housekeeping transcripts such as *GAPDH* are not modulated by eIF4E^22^. Given that most mRNAs are capped, selectivity for eIF4E-dependency arises through co-factors and RNA elements within client transcripts themselves^23, 24^. Many of these co-factors bind distal to the cap-binding site on a region of eIF4E referred to as the dorsal surface. For example, in the nucleus, eIF4E directly binds the cap on mRNAs that also contain a 4ESE element within their 3’UTR^25–28^. The large protein LRPPRC acts as a platform directly binding to the dorsal surface of eIF4E, and to the 4ESE RNA element within the target transcript ^26–28^. The eIF4E-mRNA-LRPPRC complex then binds to the nuclear exporter CRM1/XPO1 to be exported^27^. In this way, eIF4E-dependent mRNA export is selective i.e. not all capped mRNAs are targets of this process^25, 26, 29^. Similarly, eIF4E increases the translation efficiency (number of ribosomes per transcript) for specific mRNAs that contain sensitizing elements such as CERTs or long highly structured 5’UTRs^23, 30^. Many of the transcripts regulated by eIF4E encode proteins involved in biological functions related to cancer including motility^31^.

Previous studies, in solid cancer cell lines, provided a link between eIF4E motility and its capacity to drive the production of CD44 and HA^31^. This capacity had been largely attributed to the nuclear fraction of eIF4E which has multiple functions including enhanced mRNA export described above. Indeed, eIF4E drives the mRNA export, and hence cytoplasmic availability, of *CD44* and the mRNAs encoding all the enzymes involved in HA production in high-eIF4E AML and osteosarcoma cells^31^. Additionally, a few of these transcripts including *CD44* also have their translation efficiency (number of ribosomes per transcript) increased by eIF4E overexpression as well as undergo eIF4E-dependent splicing^15, 31^. Consistent with these findings, eIF4E overexpression promoted cell migration and invasion in many solid tumors and is strongly correlated with increased metastasis in patients in this setting^32–37^.

Here, we show that eIF4E promotes AML cell invasion, BM engraftment and disease progression, placing disrupted mRNA metabolism alongside dysregulated transcription and signaling as a driver of AML biology. In high-eIF4E cells, invasion depended on Ezrin, whereas the residual invasion of eIF4E-depleted cells did not. Dissecting this relationship, we found that Ezrin binds eIF4E directly through its FERM domain and associates with translation initiation factors, ribosomes and the mRNAs encoding CD44 and the HA biosynthetic pathway. To ask where translation occurs, we developed VISTA-R (<u>Vis</u>ualizing <u>T</u>ranslation <u>A</u>ctivity using <u>R</u>iboLace), which labels active ribosomes in fixed cells rather than the newly synthesized proteins that diffuse away from them within seconds. VISTA-R revealed active ribosomes in Ezrin-enriched pseudopods, which we term translationally active pseudopods (T-PODs). Ezrin and eIF4E were each required for active ribosomes to accumulate in T-PODs and for production of CD44 and HA-pathway proteins, without affecting global protein synthesis. Together, these findings reveal a direct connection between the translation and motility machineries and identify a localized, Ezrin-dependent translation program that supports AML cell movement.

## Results

### eIF4E drives AML cell migration, infiltration and disease progression

The mechanisms that control the ability of AML cells to migrate and infiltrate are not well understood, specifically the extent to which these depend upon post-transcriptional and translational control. To examine whether eIF4E activity impacted AML cell motility and infiltration, we generated two AML cell line models. For gain of function, we employed NOMO-1 AML cells since these have eIF4E levels similar to healthy donor CD34+ cells^15^ (Supplemental Figure 1A) allowing us to assess the effects of stable eIF4E overexpression pools (NOMO-1 eIF4E) relative to vector controls (NOMO-1 Vector). Here, eIF4E overexpression was sufficient to recapitulate the elevated, substantially nuclear eIF4E phenotypes typical of high-eIF4E AML primary specimens^15^. eIF4E was elevated ∼2-fold in NOMO-1 eIF4E cells versus NOMO-1 Vector^15^ which is well within the range of its elevation observed in AML patients to date^12, 13^ (Figure 1A, Supplemental Figure 1C). For loss of function, we used CRISPR to deplete eIF4E in Mono-Mac-6 (MM6) AML cells generating clonal MM6 CRISPR-4E cell lines. Parental MM6 cells have highly elevated eIF4E levels with substantial nuclear localization (Supplemental Figure1A and B) similar to those in high-eIF4E AML patients. Reduction of eIF4E in MM6 CRISPR-4E cell lines was ∼3 fold compared to CRISPR control (CRISPR-CTRL) clonal cell lines as observed by assessing multiple clones (Figure1B, Supplemental Figure 1D). Sanger sequencing confirmed that the disruption was heterozygous. Notably, complete deletion of eIF4E is lethal.

**Figure 1.**
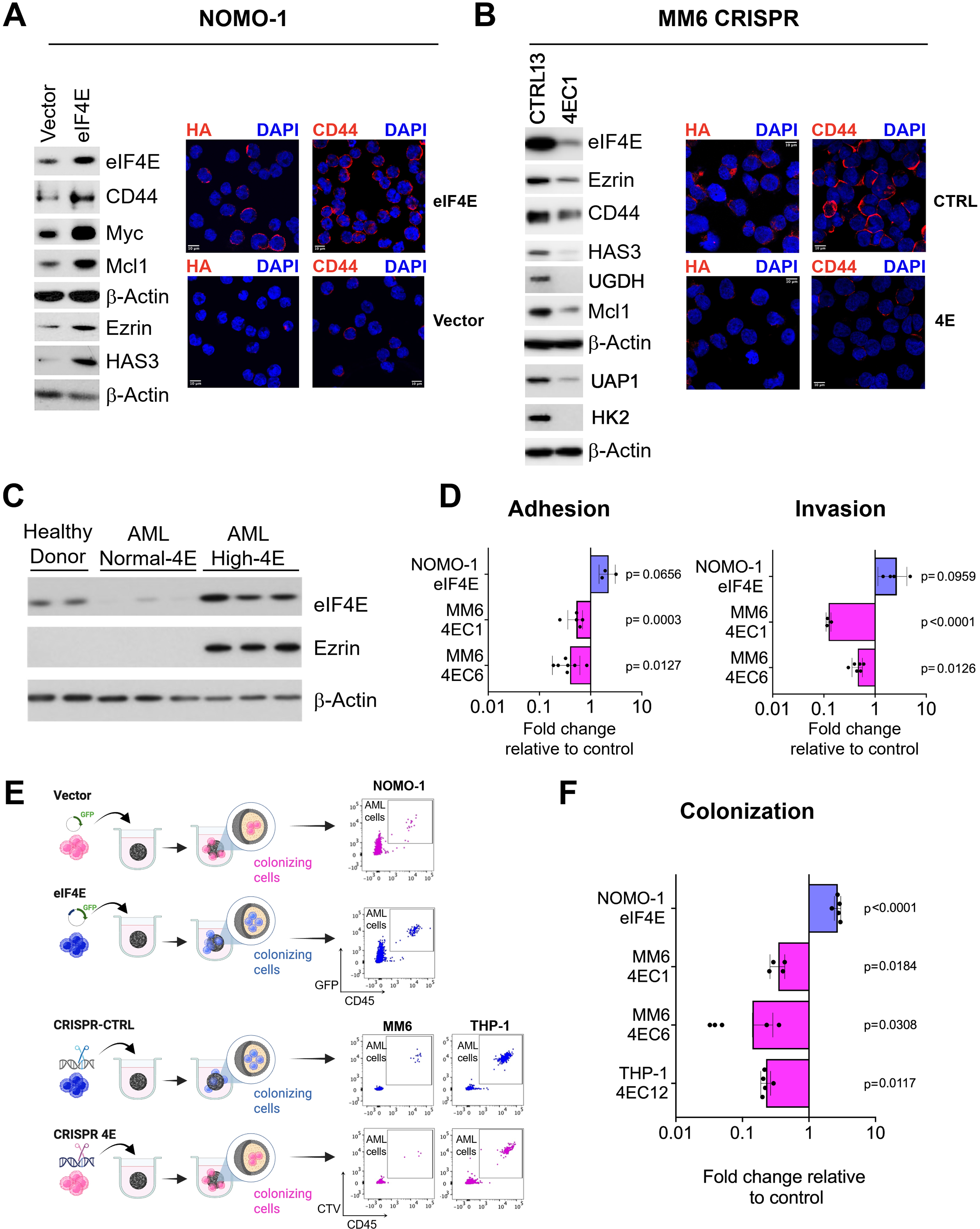
eIF4E controls cell adhesion, migration and colonization *in vitro*. **A, B.** The impact on the Ezrin-CD44-HA axis of NOMO-1 eIF4E overexpression (eIF4E) compared to vector control (A) or of MM6 CRISPR-4E cells relative to MM6 CRISPR-CTRL cells (B). **Left panels.** Western blots of eIF4E and Ezrin-CD44-HA axis. Corresponding β-Actin shown for loading controls. **Right panels**. Confocal micrographs (single section through the plane of cells) of the indicated cells stained for HA or CD44 shown in red. DAPI is shown in blue. Scale bar = 10µm. **C.** Western blot of primary specimens from AML patients (with normal or high-eIF4E) and CD34^+^ from healthy donors showing expression of eIF4E and Ezrin. β-Actin is provided as a loading control. Each lane represents a different individual. **D.** Adhesion (left) and invasion (right) capacity using HS-5 stromal cells of NOMO-1 and MM6 cell lines as a function of genetic eIF4E overexpression or CRISPR knockdown respectively. Data is presented as fold change relative to their corresponding controls. Each symbol represents an independent experiment performed with replicates. The bar represents the mean with standard deviations and p-values calculated with two tailed paired T test. **E.** Schematic representation of colonization assay into mesenchymal stromal cell spheroid model mimicking the bone marrow niche. **F.** Quantification of AML cell colonization capacity relative to their corresponding controls. Each symbol represents an experimental replicate. The bar represents the mean with standard deviations and p-values, Welch’s t test.

Given the eIF4E dependence of Ezrin-CD44-HA axis in motility in U2OS osteosarcoma cell line^31^, we assessed whether their production was also eIF4E-dependent in AML. We observed that elevation of eIF4E in NOMO-1 eIF4E cells relative to NOMO-1 Vector led to increased hyaluronic acid synthetase 3 (HAS3), CD44 and Ezrin protein levels as well as positive controls (Myc and Mcl1, known eIF4E targets^15, 31, 38, 39^) but did not impact β-Actin, which is a negative control (Figure 1A and Supplemental Figure 1C)^31^. Conversely, knockdown of eIF4E substantially reduced levels of CD44, Ezrin, HAS3 and other HA biosynthetic enzymes including UGDH, Hk2, UAP1, PGM5 in multiple clones of MM6 CRISPR-4E versus CRISPR-CTRL cell lines (Figure 1B; Supplemental Figure 1D and 1G). In AML cells, CD44 forms surface coats (Figure 1A-B) consistent with its role in HA-mediated extracellular signaling^31^. Next, we set out to monitor HA carbohydrate levels which we reasoned would be sensitive to eIF4E given the impact of eIF4E on several of the HA biosynthetic enzymes using fluorescently labeled HA-binding protein and confocal microscopy^31^. Here, HA levels were substantially elevated in NOMO-1 eIF4E cells versus NOMO-1 Vector; and conversely, HA levels were reduced in MM6 CRISPR-4E versus CRISPR-CTRL cell lines (Figure 1A-B). HA coated the cells consistent with our previous studies that HA was both elevated and coated leukemic cells in primary high-eIF4E AML patient specimens relative to CD34+ cells from healthy donors as well as in parental MM6 cells^31^. Importantly, treatment with hyaluronidase (HAse) eliminated the HA signal indicating that the staining was specific (Supplemental Figure 1E). Finally, we observed that Ezrin is highly elevated in primary high-eIF4E AML specimens relative to normal-eIF4E AML patients’ specimens or CD34+ cells from healthy donors (Figure 1C), consistent with these having elevated eIF4E and HA^31^; β-Actin is shown as a loading control. Thus, consistent with nuclear eIF4E RNA immunoprecipitation (RIP) and mRNA export data from U2OS cells^31^, eIF4E drives protein production of Ezrin as well as of HAS3 and CD44 in AML cells.

Since eIF4E modulates production of the Ezrin-CD44-HA axis in AML cells (Figure 1A-B), we investigated whether eIF4E played a role in AML motility by performing *in vitro* adhesion and invasion assays. For these assays, we employed the HS-5 human cell line as it recapitulates the major features of primary human BM mesenchymal stromal cells^40^. For adhesion assays, AML cells pre-stained with a fluorescent dye [DiIC12(3)] were incubated with a monolayer of HS-5 cells for 1 hour, washed and the number of cells retained on the stroma was quantified^31^. For invasion, pre-stained AML cells were loaded into the upper compartment of a Boyden chamber with HS-5 coated inserts and the C-C motif chemokine ligand 2 (CCL2, also known as monocyte attractant protein 1, MCP-1) in the lower chamber as a chemoattractant and incubated for 24 hours. Experiments with pools of stable NOMO-1 eIF4E and NOMO-1 Vector cells were conducted in biological replicates and different clones for MM6 CRISPR (4E and CTRL) cell lines were used. We observed that NOMO-1 eIF4E cells had ∼2-fold increased adhesion relative to NOMO-1 Vector, and ∼2-fold increased invasion through HS-5 stroma (Figure 1D). Conversely, MM6 CRISPR-4E cells had ∼2-fold reduced adhesion and ∼2-fold decreased invasion versus CRISPR-CTRL cells (Figure 1D). Plating controls were used to ensure equal loading of cells in both adhesion and invasion experiments (see Materials and Methods). These AML cell lines double every ∼36-60 hours and thus proliferative effects do not contribute to these results. Furthermore, there were no detectable differences in proliferation or viability for NOMO-1 eIF4E versus NOMO-1 Vector or MM6 CRISPR-4E over MM6 CRISPR-CTRL indicating that observed changes in motility are not due to differences in proliferation or survival (data not shown). In all, these studies reveal a major role for eIF4E in AML cell migration *in vitro*.

To build on these findings, we explored the eIF4E-dependency of AML infiltration in a system that mimics specialized niche structures and cell-to-cell intercommunication known to drive malignant hematopoiesis^41^. For this purpose, we used an *in vitro* functional 3D hematopoietic-mesenchymal co-culture system (spheroid) consisting of HS-5 and AML cells. Leukemia cells with infiltrating/colonizing properties are scored based on their localization inside the 3D structure (Figure 1E). Thus, using this system, we examined two different clones of MM6 CRISPR-4E cells which had between 3- and 7-fold reduced capacity for colonization on day 1 compared to two MM6 CRISPR-CTRL clonal cell lines (Figure 1F). We extended this into another high-eIF4E AML cell line THP-1 which was previously characterized by a >3-fold level of intrinsic eIF4E elevation relative to healthy donor specimens^14^. We generated THP-1 CRISPR-4E (THP-1 4EC12) and CRISPR-CTRL cells (THP-1 CTRL10) as for MM6 cells. Western blots confirmed knockdown of eIF4E corresponded to reduction in CD44, Ezrin and Myc in THP-1 CRISPR-4E cells (Supplemental Figure 1H). These data are consistent with observations in CRISPR-4E THP-1 cells where we observed a 5-fold decrease in colonization relative to THP-1 CRISPR-CTRL cells and consistently, we found that NOMO-1 eIF4E cells had a 2.9-fold increase in colonization capacity relative to vector controls. Taken together, we demonstrated that eIF4E modulation impacts the ability of leukemia cells to invade and colonize the spheroid which represents the BM niche. We also generated tools for *in vivo* study of the impacts of eIF4E in the pathogenesis of AML.

As we previously showed that colonizing cells in this model reflect cells with increased engraftment capacity in xenografts^41^, we monitored the impact of eIF4E depletion on the engraftment of AML cells to the BM niche employing the two MM6 and one THP-1 clonal cell lines for CRISPR-4E and CRISPR-CTRL using xenotransplantation into immunodeficient mice. We evaluated AML cell engraftment in BM at a timepoint when the control animals were expected to succumb to the disease (Figure 2A and Supplemental Figure 2A and 3A). We found a significant decrease in engraftment in mice transplanted with MM6 CRISPR-4E (clone 4EC6) cells compared to MM6 CRISPR-CTRL (clone CTRL7) cells [% of human cells in BM (mean ± SE): 10.13 ± 2.78 (n=10) vs. 56.53 ± 4.98 (n=10), respectively, p<0.0001] (Figure 2B). These results were corroborated on a different set of clonal cell lines, MM6 CRISPR-4E (clone 4EC1) cells compared to MM6 CRISPR-CTRL (clone CTRL13) cells [% of human cells in BM (mean ± SE): 0.26 ± 0.14 (n=5) vs. 5.20 ± 1.51 (n=5), respectively, p=0.0079, Supplemental Figure 2B]. Importantly, these observations were validated using another set of CRISPR-4E and CRISPR-CTRL in THP-1 cells. THP-1 cells needed longer time to engraft but still caused disease in the animals. Thus, when we first evaluated engraftment at day 30 prior to any symptoms of disease, we found that 80% of the animals transplanted with THP-1 CRISPR-CTRL (clone CTRL10) had detectable human cells in the BM, in stark contrast with 10% of the THP-1 CRISPR-4E (clone 4EC12, Supplemental Figure 3B). Next, we waited until the control animals started to display signs of disease, at this point day 56. As we sacrificed 3 succumbing THP-1 CRISPR-CTRL mice, we took 3 CRISPR-4E mice to compare the engraftment levels in BM, and spleen, confirming a higher engraftment of human cells in the CTRL mice when compared with CRISPR-4E (Supplemental Figure 3C). At day 59, all CTRL animals had to be sacrificed due to disease progression, we isolated the BM from these animals to compare with BM aspirates from all CRISPR-4E surviving mice (with no signs of disease). We again observed a significant decrease in engraftment in mice transplanted with THP-1 CRISPR-4E (clone 4EC12) cells compared to THP-1 CRISPR-CTRL (clone CTRL10) cells: [% of human cells in BM (mean ± SE): 2.5 ± 0.87 (n=6) vs. 8.97 ± 2.43 (n=6), respectively, p=0.0450] (Figure 2B).

**Figure 2.**
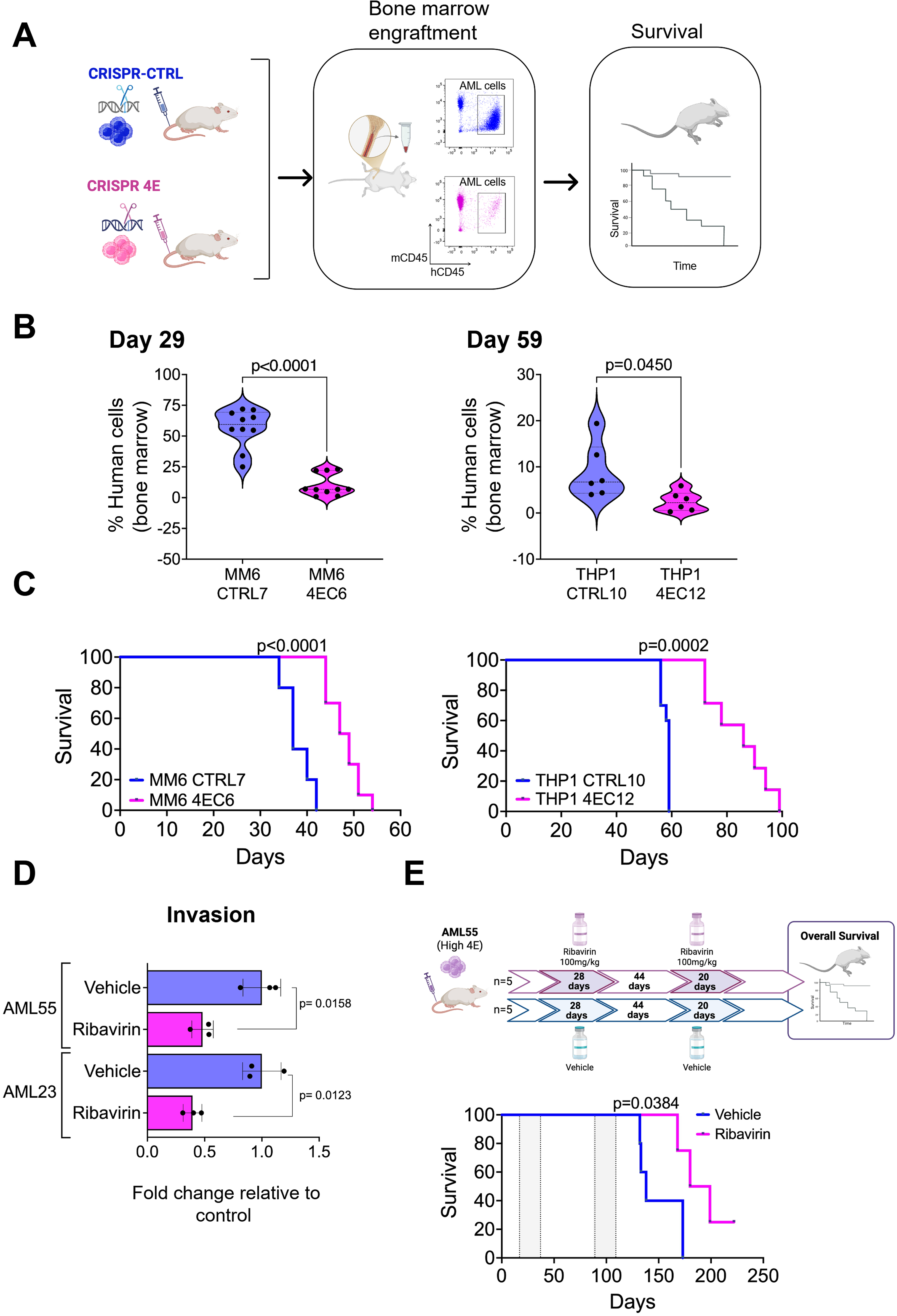
Depletion of eIF4E reduces engraftment and improves overall survival in AML cell line-derived xenograft (CDX) mouse models. **A**. Schematic representation of experimental design using CDX mouse models. **B.** Percent human cells in the bone marrow at the indicated timepoint assessing the AML engraftment capacity for the indicated cell lines, each dot represents an animal. Violin plot representing the median and quartiles, p-values calculated with unpaired two tailed t test and Welch’s t test. **C.** Kaplan–Meier curves comparing overall survival of mice transplanted with the indicated cell lines. Log-rank (Mantel-Cox) test applied. **D.** invasion capacity through HS-5 stromal cells of the indicated primary AML samples after exposure to Ribavirin or vehicle control. Data is presented as fold change relative to vehicle control. Each symbol is an independent experiment performed with replicates. The bar represents the mean with standard deviations and p-values calculated with Welch’s t test (two tailed). **E.** Pharmacological targeting of eIF4E with Ribavirin reduces leukemia burden and improves overall survival in an AML patient-derived xenograft (PDX) mouse model. Diagram illustrating the treatment regimen (top panel). Kaplan–Meier curves comparing overall survival of PDX mice with the indicated treatments. Log-rank (Mantel-Cox) test applied (bottom).

Consistently, decreased engraftment resulted in a significant improvement in overall survival (OS) in animals transplanted with CRISPR-4E cells. Specifically, for MM6 cells OS was 48 days for CRISPR-4E (4EC6 n=10) vs 37 days for CTRL (CTRL7 n=10), p<0.0001 (Figure 2C). Similarly, THP-1 CRISPR-4E (THP-1 4EC12, n=7) had an improved OS of 86 days compared with 59 days for THP-1 CRISPR-CTRL cells (THP-1 CTRL10; n=10), p=0.0002 (Figure 2C). The additional MM6 clone showed an OS of 51 days for CRISPR-4E (4EC1; n=5) vs 32.5 days for CTRL (CTRL13; n=4), p=0.0040 (Supplemental Figure 2C). Of note, all animals (n=10) injected with THP-1 CRISPR-CTRL cells succumbed with extramedullary tumors (abdomen) while 5 animals from the THP-1 CRISPR-4E cohort presented small abdominal tumors (Supplemental Figure 3D), suggesting that the deficiency in eIF4E may have also prevented the cells from migrating to form abdominal tumors. Finally, we isolated human leukemic cells from the mice, and these cells retained their reduced levels of eIF4E (Supplemental Figures 2D and 3E). In summary, when transplanted into mice, cells with reduced eIF4E levels which correspond to reduced levels of Ezrin, CD44, and HA (Figure 1A-B) were characterized by reduced engraftment with concomitant increased overall survival in these animals, demonstrating a role for eIF4E in controlling these processes (Figures 1 and 2, Supplemental Figures 2 and 3).

### Pharmacological targeting of eIF4E and motility

In addition to the genetic manipulation of eIF4E, we studied pharmacological inhibition of eIF4E to test the impact on OS using primary AML cells as patient derived xenografts (PDXs). To date, the only therapy that elicited eIF4E-targeting corresponding to objective clinical improvement in patients utilizes the antiviral drug ribavirin which can act as a m^7^G-cap competitor and thus eIF4E inhibitor as shown by NMR and other biophysical techniques^42–44^. Ribavirin inhibits eIF4E’s activities in mRNA export, translation, and oncogenic transformation in cell lines, animal models and patients as shown by over 30 groups^14, 33, 42, 43, 45–50^. These studies include demonstrations that eIF4E overexpression can rescue cells from ribavirin treatment and, moreover, that ³H ribavirin binds to eIF4E in intact cells.^32, 42, 48–50^. Three clinical trials using ribavirin alone or in combination to treat refractory/relapsed high-eIF4E AML patients led to objective clinical responses including complete remissions correlating with molecular targeting of eIF4E^12, 13, 51^ and promising results were observed in oral cancer^52^. While there was some evidence that ribavirin may have very weak IMPDH inhibitory activity in cells that accumulate the rare ribavirin metabolite RMP, more recent studies demonstrate that ribavirin synergizes with FDA approved IMPDH inhibitors and that ribavirin does not generally act as an IMPDH inhibitor^53^. In all, while ribavirin like all small molecules can have additional activities, it inhibits eIF4E.

Similar to genetic targeting in parental MM6 cells, Ezrin and HAS3 protein levels were reduced by ribavirin as were positive controls Myc and Mcl1 while, the negative control, β-Actin was unaffected (Supplemental Figure 1F). We compared the invasion capacity of high-eIF4E AML primary samples (AML55 and AML23) after ribavirin treatment *ex vivo*. We observed that both specimens had approximately 2-to3-fold reduction in invasion after ribavirin treatment relative to vehicle treated cells (p= 0.0158 and 0.0123 respectively) (Figure 2D). Thus, ribavirin treatment reduced AML55 and AML23 invasion nearing that to healthy donors (Figure 2D). Moreover, AML55, AML23 and another high-eIF4E sample AML13 had ∼2-to 7-fold increased invasion relative to CD34+ cells from healthy donors (Supplemental Figure 4A).

To extend this into an *in vivo* model, we established AML55-PDX mice which received two cycles of ribavirin (100mg/kg) (Figure 2E upper panel). Peripheral blood (PB) samples were obtained after the end of the second cycle (day111) where we found that animals treated with ribavirin had significantly less AML cells than mice treated with vehicle control (0.9% ± 0.24; n=5, vs. 7.5% ± 3.8; n=5, respectively, p=0.0079) (Supplemental Figure 4A and B). Approximately a week later, the fast progression of AML is observed in the Vehicle control (saline) mice when compared with the ribavirin treated group (44.9% ± 44.9; n=5, vs. 2.4% ± 1.65; n=5, respectively, p=0.0079) (Supplemental Figure 4B). Importantly, mice that received ribavirin survived ∼51 days longer than the control animals treated with saline control (ribavirin cohort median survival=189 days vs saline=138 days; p=0.0384). Altogether, we demonstrated that the genetic depletion or pharmacological inhibition of eIF4E impairs engraftment within the BM niche and improves overall survival in AML *in vitro* and *in vivo* models and further support the importance of eIF4E for AML cell motility.

### A novel interplay between the motility and translation machineries

Given the role of eIF4E in motility shown above and the established relevance of the Ezrin-CD44-HA circuit in AML, we first assessed whether AML cell motility required Ezrin. Here, we monitored the impact of genetic reduction on adhesion and invasion through HS-5 BM stroma cells using siRNA to *EZR* in MM6 cells and compared to siRNA to luciferase (siLUC; which served as negative control). siRNA to *EIF4E* was used for comparison. RNAi reduced Ezrin levels by >10-fold relative to siLUC as observed by western blot with no significant impact on eIF4E protein levels (0.94±0.12; p-value=0.08; Figure 3A, Supplemental Figure 5A). Decreased Ezrin levels reduced adhesion by ∼2-fold and invasion by ∼1.5-fold relative to siLUC controls (Figure 3B). By comparison, reduced eIF4E levels after siRNA (0.27±0.08) led to decreased levels of Ezrin and CD44 corresponding to a ∼2-fold reduction of adhesion and invasion relative to siLUC control (Supplemental Figure 5A), similar to the decreased levels of Ezrin and CD44 in the CRISPR-4E versus CRISPR-CTRL cells (Figure 1B; Supplemental Figure 1D) and subsequent ∼2-fold reduction of adhesion and invasion in CRISPR-4E versus CRISPR-CTRL MM6 cells (Figure 1D). We note that siEIF4E caused the anticipated reduction in CD44 and HA biosynthetic enzymes similar to CRISPR-4E (Supplemental Figure 1G). Interestingly, Ezrin knockdown caused similar changes to CD44 and HA pathway enzymes which are explored below. In all, Ezrin is required for AML cell motility garnering similar effects to eIF4E.

**Figure 3.**
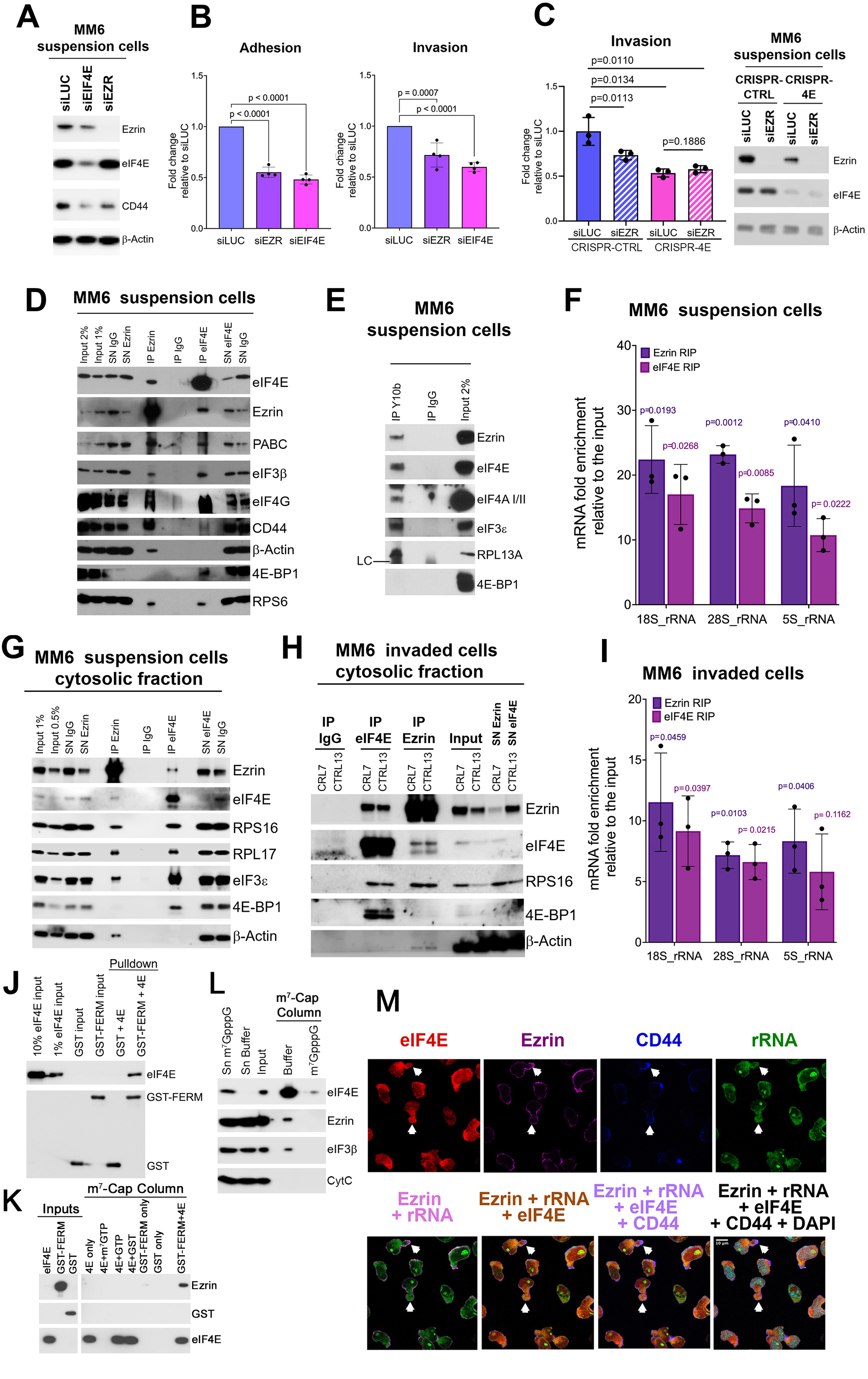
Ezrin and eIF4E functionally and physically interact. **A.** Western blot of total cell lysates from MM6 cells grown in suspension demonstrated knockdown of Ezrin (siEZR) or eIF4E (siEIF4E) compared to RNAi to luciferase (siLUC) used as a negative control. β-Actin is provided as a loading control. Other proteins of the Ezrin-CD44-HA axis are also shown. Quantification for these is shown in Supplemental Figure 5A with 3-6 biological replicates for each protein. **B.** Adhesion and invasion capacity of MM6 cells onto/through HS-5 bone marrow stroma. Fold change relative to siLUC is shown. Each symbol represents a biological replicate performed independently with technical replicates. Bars represent the mean, shown with standard deviations and p-values (one-way ANOVA). **C.** Left panel: Ezrin knockdown effect on invasion capacity of MM6 CRISPR-4E cells relative to siLUC controls cells through HS-5 bone marrow stroma. Fold change relative to siLUC is shown. Each symbol represents a biological replicate performed independently with replicates. Right panel: Western blot of total cell lysates from MM6 cells grown in suspension demonstrating knockdown of Ezrin (siEZR). **D.** Western blot of eIF4E and Ezrin immunoprecipitations (IPs) using total cell lysates from MM6 cells in suspension. SN, supernatant after immunoprecipitation, IgG, negative control. Representative of three biological replicates. IPs of total cell lysates from THP-1 cells in suspension are provided in Supplemental Figure 5B. **E.** IPs from MM6 cells in suspension using the rRNA antibody Y10b. LC indicates antibody light chain. Representative of three biological replicates. **F.** RNA immunoprecipitations (RIPs) from MM6 total cell lysates grown in suspension performed using anti-Ezrin (Ezrin RIP) or anti-eIF4E (eIF4E RIP) antibodies. Data are from RT-qPCR and represented relative to input. Each symbol represents a biological replicate performed independently with technical replicates. Bars represent the mean, shown with standard deviations and p-values (two tailed Welch’s t test). **G** and **H.** Western blots of eIF4E and Ezrin IPs from the cytoplasmic fractions of MM6 cells in suspension (G) or after invasion through HS-5 bone marrow stroma (invaded) (H). Fractionation controls are provided in Supplemental Figure 5E. **I.** RIPs from MM6 cytoplasmic fraction from invaded cells using anti-eIF4E (eIF4E RIP) or anti-Ezrin (Ezrin RIP) antibodies. Data are from RT-qPCR represented relative to input. Each symbol represents a biological replicate performed independently with triplicates. Bars represent the mean, shown with standard deviations and p-values (two tailed Welch’s t test). **J**. WB analysis of a GST pulldown assay using purified Ezrin GST-FERM and eIF4E, demonstrating a direct interaction between Ezrin and eIF4E. **K**. WB analysis of m7G-cap chromatography using purified proteins, showing that Ezrin binds to the m7G-cap column only in the presence of eIF4E. **L**. WB analysis of m7G-cap binding using cytoplasmic lysates from MM6 cells, showing binding of endogenous Ezrin to m7G-cap beads. **M**. Immunofluorescence and confocal microscopy demonstrating eIF4E, Ezrin, CD44 and rRNA are localized to the same pseudopods (white arrows). All confocal micrographs represent a single section through the plane of the cell. Scale bar = 10 µm.

While Ezrin plays wide roles in motility, we assessed whether high-eIF4E AML cells were more sensitive to Ezrin loss than cells with normal eIF4E levels. MM6 CRISPR-4E cell lines retain approximately half the invasive capacity of MM6-CTRL cells with high eIF4E expression and therefore still exhibit measurable invasion capacity (Figure 1D). Here we observed that Ezrin knockdown had no impact on the invasion capacity of MM6 CRISPR-4E cells relative to siLUC controls (Figure 3C) i.e. that the invasion in these was not sensitive to Ezrin reduction. Conversely, reduction of Ezrin decreased invasion about 30-40% of MM6 CTRL cells compared to siLUC controls (Figure 3C). Thus, high-eIF4E AML cells rely on Ezrin more for motility than cells with reduced eIF4E levels. This suggests there is more of a functional interplay between Ezrin and eIF4E in motility in the high-eIF4E context.

Interestingly, we noted that knockdown of both Ezrin and eIF4E reduced levels of CD44 protein and correspondingly impaired eIF4E-dependent motility without affecting eIF4E protein levels (Figure 3A, Supplemental Figure 5A). Given Ezrin’s localization is limited to the cytoplasm, these observations led us to speculate that Ezrin played an unanticipated physical role in translation. To address this, we investigated whether there were physical interactions between eIF4E and Ezrin using immunoprecipitation of endogenous factors (Figure 3D-J; Supplemental Figure 5B). We took two approaches: 1. MM6 and THP-1 cells were crosslinked immediately prior to collection to avoid reassortment during processing and immunoprecipitation and 2. native immunoprecipitations. Using either strategy, we observed that endogenous eIF4E immunoprecipitated with Ezrin in total cell lysates from both cell lines as well as in cytoplasmic fractions in MM6 cells (Figure 3D, G and Supplemental Figure 5B and C). Conversely, endogenous Ezrin immunoprecipitated with eIF4E as well as CD44 and Actin in total cell lysates as well as from cytoplasmic fractions (Figure 3D, G, Supplemental Figure 5B and 5C). Actin served as a negative control for eIF4E. Interestingly, eIF4E also immunoprecipitated with CD44, which is known to directly bind to Ezrin at the plasma membrane^7^ (Figure 3D, Supplementary Figure 5B). eIF4E immunoprecipitated with its negative regulator 4E-BP1 consistent with it existing as a mixed population of active and inactive forms, as described in the literature^54^. Notably, 4E-BP1 did not associate with Ezrin suggesting that Ezrin interacts with the active form of eIF4E. Factors were not observed in the IgG negative controls. Next, we explored whether the interactions between Ezrin and eIF4E were bridged by RNA by treating Ezrin IPs conducted under native conditions with a combination of RNAse I, A and T2 to reliably degrade structured and unstructured RNA forms. RNAse treatment did not significantly reduce the Ezrin–eIF4E interaction (Supplementary Figure 5D).

The RNA independence for the endogenous eIF4E-Ezrin interaction suggested the possibility that Ezrin and eIF4E directly interacted. Purified eIF4E and the FERM domain of Ezrin fused to GST (GST-FERM) were produced in bacteria, purified and interactions assessed using GST pulldown assays. Here, eIF4E, in the absence of the cap, directly interacted with Ezrin GST-FERM but not the GST negative control (Figure 3J). We next examined whether Ezrin GST-FERM can bind to eIF4E-m^7^G cap complexes. For this purpose, we used cap chromatography where an m^7^G-cap analog is immobilized on agarose beads and incubated these purified eIF4E, Ezrin GST-FERM or GST proteins. We observed that eIF4E bound the beads as expected and that Ezrin GST-FERM bound to the m^7^G-cap beads in the presence of eIF4E but not in its absence. GST did not bind the beads even in the presence of eIF4E. Thus, Ezrin binds to eIF4E-cap complexes (Figure 3K). Importantly, addition of the m^7^cap analog completely abolished the binding of eIF4E to m^7^G-cap beads demonstrating this is a cap-dependent interaction (Figure 3K). Consistent with our *in vitro* study, we found that endogenous eIF4E bound to the m^7^G-cap beads after incubation of MM6 cytoplasmic lysates as did Ezrin and the positive control eIF3β (Figure 3L). Addition of a m^7^G-cap analog abrogated the binding of eIF4E and thus eIF3β and Ezrin. In all, eIF4E and Ezrin directly interact and moreover, Ezrin-eIF4E-capped mRNA complexes exist.

Given these findings, we predicted that Ezrin would co-exist in complexes with translation factors mediated through eIF4E (Figure 3D-I; Supplemental Figure 5B-D). Consistent with our *in vitro* studies, endogenous Ezrin immunoprecipitated with several key components of the translation initiation factors including eIF4AI/II, eIF4G, eIF3 components (3β and 3ε), and the ribosomal small and large subunit proteins RPS6, RPS16 and RPL17. Endogenous eIF4E immunoprecipitated with these factors as expected. We also observed these interactions in THP-1 cells as well (Supplemental Figure 5B). To further solidify Ezrin’s physical association with the translation machinery, we assessed if Ezrin was found in the same complexes as ribosomal rRNAs. Thus, we carried out RNA immunoprecipitation (RIP) and RT-qPCR monitoring interactions with ribosomal rRNAs (5S, 18S and 28S rRNAs) from total cell lysates of MM6 cells. Indeed, Ezrin RIPs were characterized by an 18-to 22-fold enrichment in rRNAs over input in MM6 cells (Figure 3F). For comparison, eIF4E RIPs were 10-to17-fold enriched for these rRNAs. Differences in fold changes between Ezrin and eIF4E RIPs likely arise due to differences in avidity of the antibodies and cannot be used to establish different affinities. To independently validate that Ezrin and ribosome were in the same complexes, we employed the Y10b antibody to rRNAs^55^. Using crosslinked total cell lysates, we observed that Y10b immunoprecipitated Ezrin as well as positive controls eIF4E and the translation machinery but not the 4E-BP1 (Figure 3E). In all, Ezrin co-exists in complexes with the translation machinery likely reliant on its direct interaction with eIF4E.

Given the considerable structural rearrangements in cells upon invasion including the formation of pseudopods, we monitored interactions of these factors in invaded cells. After invasion, we fixed MM6 cells from the lower compartment of the Boyden chamber, collected these and monitored whether Ezrin interactions with the translation machinery were preserved. With this material, we conducted immunoprecipitation experiments of cytoplasmic fractions from invaded cells (Figure 3G and I). We note that there is substantial reduction in invaded cell number relative to cells grown in liquid culture yielding only about 100 µg of material which limited the extent of immunoprecipitation analyses. We observed that eIF4E and Ezrin co-immunoprecipitated with the ribosomal RPS16 protein. Ezrin did not immunoprecipitate with 4E-BP1 while eIF4E did, consistent with findings in cells grown in suspension. Moreover, both Ezrin and eIF4E immunoprecipitated with rRNAs with 5-16-fold enrichment relative to input (Figure 3I). Thus, Ezrin also co-exists in the same complexes as the translation machinery and ribosomes in invaded cells. MEK, CalR and Nopp140, H3K27A served as cytoplasmic and nuclear markers respectively to show the quality of the fractionation (Supplemental Figure 5E). Given Ezrin is restricted to pseudopods in invaded cells while eIF4E is found throughout the cytosol and in pseudopods, we expected eIF4E to bind more mRNAs and rRNAs. We note that the fold-enrichment between Ezrin and eIF4E RIP or IP experiments likely also reflect differences in antibody avidity restricting any interpretations of differences in binding affinity. In all, immunoprecipitation of Ezrin with the translation machinery is preserved in invaded cells. Here, we show that Ezrin directly binds eIF4E and co-exists in complexes with the translation machinery in multiple cellular contexts suggesting that Ezrin plays an important role in specialized translation likely to produce factors required for motility.

### Pseudopods in motile AML cells contain eIF4E and rRNAs

Given the relevance of pseudopods to AML cell motility and the association of CD44 and Ezrin with these structures^56,57^, we set out to assess if these could be sites of novel form of eIF4E-Ezrin dependent localized translation. To assess the best conditions to monitor pseudopods, we set out to quantify the difference in pseudopod numbers per AML cell before and after HS-5 BM stroma invasion. We observed most invaded cells (∼80%) had at least one pseudopod, while ∼35% of suspension cells (grown under regular conditions, and without going into invasion process) exhibited pseudopods (Supplemental Figure 5F). Given the higher prevalence of pseudopods in invaded cells, we focused on these for the remaining studies unless otherwise noted.

Next, we monitored the spatial localization of eIF4E and ribosomal rRNAs relative to Ezrin and CD44 employing immunofluorescence and confocal microscopy with Z-stacks (Figure 3M). To capture invaded cells, cells in the lower compartment of the Boyden chamber after invasion through HS-5 cells were fixed with paraformaldehyde (PFA) *in situ*, collected and then underwent processing for immunofluorescence to assess the localization of eIF4E, Ezrin, CD44, rRNA (Y10b antibody) and DAPI as a nuclear marker (Figure 3M). To ensure there was no physical or optical interference, each wavelength was collected independently, and single staining controls showed no reorganization or bleed through of signal between single and multiple staining experiments (data not shown). Confocal settings provide a resolution of ∼200 nm (see Materials and Methods). In MM6 cells in suspension, eIF4E is found in nuclear bodies as well as in the cytoplasm as reported^58–61^ (Supplemental Figures 1B). In invaded MM6 cells a substantial fraction of eIF4E and rRNA are also, interestingly, located in pseudopods where the majority of Ezrin and CD44 are found (Figure 3M). We observed a highly similar pattern for eIF4E localization in invaded relative to suspension cells with both characterized by nuclear eIF4E and cytoplasmic eIF4E, as well as an eIF4E-enrichment in pseudopods in these cells (Figure 3M). As expected, rRNAs were also found in the nucleolus. Thus, consistent with the immunoprecipitation data (Figure 3 D-I), we observed that Ezrin was localized to the pseudopods containing eIF4E, rRNA and CD44 (Figure 3M).

### Active ribosomes are found in pseudopods

Given the localization of eIF4E, ribosomal proteins, and Ezrin we sought to ascertain whether pseudopods contained active ribosomes, i.e. sites of translation, or if they act as storage sites for translation factors and thus are sites of sequestration. To this end, we developed a method to directly mark active ribosomes *in situ*. Several limitations in previous methods used to monitor *in situ* translation prompted this step. Foremost is that proteins diffuse from the site of protein synthesis within seconds^62^. Thus, the use of labeled amino acids or puromycin (or its analogs) to mark sites of translation is problematic since these rely on the production of newly synthesized labels such as puromycilated proteins to act as proxies for translation but these newly synthesized proteins rapidly diffuse from ribosomes^62^ (Figure 4A, left panel). This clearly confounds determination of the actual sites of translation. To overcome this, we developed a new assay: <u>Vis</u>ualizing <u>T</u>ranslation <u>A</u>ctivity using <u>R</u>iboLace (VISTA-R). VISTA-R is an *in situ* translation assay which employs the biotinylated puromycin analog known as 3P molecule for RiboLace, to directly detect active ribosomes^63^ rather than monitoring newly synthesized puromycilated proteins (Figure 4A right panel, Supplemental Figure 6A). Puromycin and its analogs enter the empty A-site of the active ribosome mimicking a tRNA, which leads to premature chain termination, release of a puromycilated peptide, and ribosomal dissociation^63^. To overcome these issues in VISTA-R, cells were fixed with PFA to freeze ribosomes prior to incubation with 3P molecule for RiboLace so that the ribosomes themselves are marked rather than visualizing newly synthesized proteins. 3P (RiboLace) was subsequently detected by streptavidin-fluorophore conjugate and confocal microscopy. In this scenario, we are not monitoring the incorporation of the 3P molecule into the peptide chain but rather directly labelling active ribosomes.

**Figure 4.**
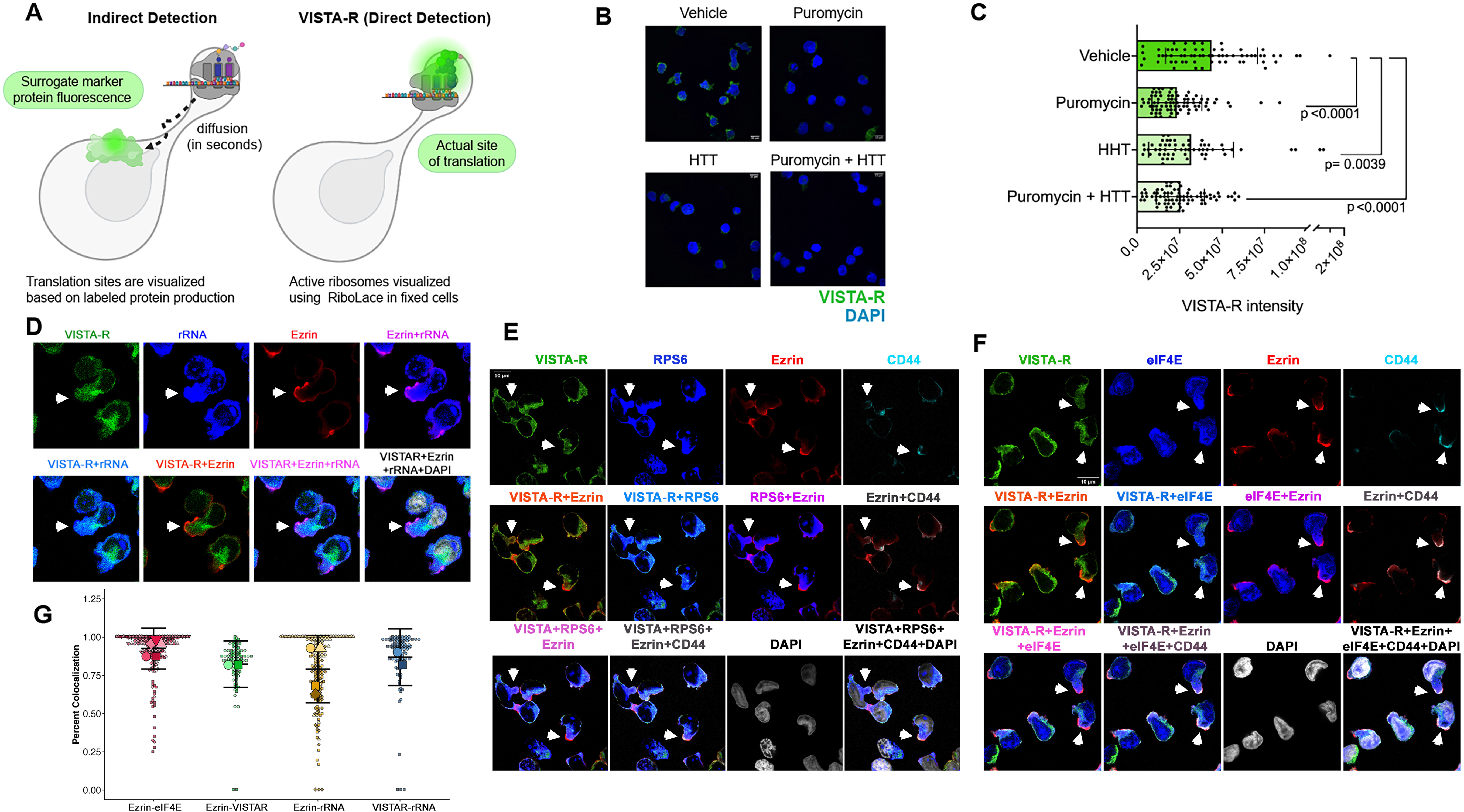
Active ribosomes and translation machinery are observed in T-PODs. **A**. Schematic comparing previous strategies to monitor in situ translation based on surrogate markers of translation such as labeled amino acids (left), and our novel method for direct measurement of translation, VISTA-R (right). **B.** VISTA-R signals in MM6 cells are directly related to translation activity since these are abrogated upon addition of translation inhibitors. Puromycin and/or HHT were added prior to the fixation step in VISTA-R. Representative confocal micrograph of VISTA-R signal in vehicle control, puromycin, HHT or combination treated cells. DAPI is shown in blue. Scale bar = 10 µm. **C.** Quantitation showing means, standard deviations of the mean, data points from each measurement, p-values calculated with one way ANOVA test from over 60 cells from two independent experiments. **D.** T-PODs (as shown by arrows) contain active ribosomes as seen by VISTA-R (green). VISTA-R and rRNA (red), Ezrin (blue) overlay in these T-PODs. **E.** Same as D but demonstrating VISTA-R (green), Ezrin (magenta) and RPS6 (red) and CD44 (turquoise) are present in the same T-PODs (see arrows). **F.** Same as D-E but demonstrating that VISTA-R (green), eIF4E (blue) and Ezrin (red) and CD44 (cyan) are present in the same T-PODs (see arrows). Colocalization of the green and red channels appears in orange, while colocalization of the blue and red channels appears in pink. All confocal micrographs represent a single section through the plane of the cell. Scale bars = 10 µm. **G.** Analysis of Ezrin colocalization with translation machinery factors using the Imaris Overlapped Volume Ratio to Surfaces metric. Data are presented as Superplots. Each small symbol represents an individual cell; different shapes represent independent experiments, with larger symbols representing the mean for each individual experiment. Horizontal lines indicate the median ± standard error across all independent experiments.

Using VISTA-R and confocal microscopy, we observed signals throughout most of the cytoplasm, and importantly, there was no nuclear signal detected consistent with the absence of diffusion using this strategy (Figure 4B, D-F). To validate VISTA-R staining was marking active ribosomes, we monitored the impacts of translation inhibitors on the VISTA-R signal. For example, we first inhibited translation elongation using puromycin (30 minutes at 250µg/mL) to allow the ribosomes to dissociate and then fixed the cells in PFA after which we followed the VISTA-R protocol and quantified signals in Imaris software. Here, we monitored signal across the entire cell volume and observed a ∼2-fold reduction in signal (green) upon puromycin treatment consistent with the VISTA-R method detecting active translation (Figure 4C). We observed similar results using the translation inhibitor homoharringtonine (HHT) (30 minutes, 10µM) which also binds the A-site but stalls translation initiation^64^. Combination of HHT and puromycin for 30 minutes also reduced the VISTA-R signal by ∼2-fold. These are strong reductions in VISTA-R signals considering these treatments were only for 30 minutes (Figure 4B and C). We note that complete abrogation of the signal would not be expected given that there is competition between reinitiation of translation and reassembly of the ribosome after inhibition. For method development, we note that we used suspension rather than invaded cells given the lower number of cells that could be obtained from the invasion experiments (Figure 4B). In all, VISTA-R detects active ribosomes *in situ*.

We combined VISTA-R with immunofluorescence to monitor if Ezrin-positive pseudopods were sites of translation using spinning disk confocal microscopy (Figure 4D-F). In invaded cells, pseudopods contained VISTA-R and rRNAs which overlapped (in orange, see arrows, Figure 4D) again indicative of active ribosomes in these structures and further confirming VISTA-R is marking these structures. Ezrin, eIF4E, VISTA-R, RPS6, and CD44 were also found in the same pseudopods where they partially overlapped (Figure 4E in white, see arrows) consistent with the above immunoprecipitation data (Figure 3D-I). With a focus on Ezrin function, we quantified the extent that Ezrin overlapped with target signals using the Imaris overlapped volume ratio to surfaces metric. This analysis revealed extensive colocalization of Ezrin with the translation machinery: 0.925 of Ezrin overlapped with eIF4E, 0.825 overlap of Ezrin with VISTA-R and 0.791 overlap of Ezrin with rRNA (Figure 4G), and consistent with the above controls, the overlap of VISTA-R with rRNA was 0.868. While there is certainly eIF4E, VISTA-R and rRNA signals beyond Ezrin given their wider distribution, our results show that much of the Ezrin protein is localized with the translation machinery. We note that the spatial localization of these factors within pseudopods is not uniform suggestive of compartmentalization within these structures. We observed that Ezrin is often most highly enriched at the tip of the pseudopods but is also found within their interior (Figure 4D-F, arrows). The overlap of Ezrin with eIF4E, rRNAs and active ribosomes is most prominent at the tips of these structures (in white). To investigate the clinical relevance of pseudopods we examined a primary high-eIF4E AML patient specimen (AML23) which displayed similar active ribosome-positive, Ezrin-positive pseudopods (Supplemental Figure 6B) as well as increased invasion over healthy controls (Supp Figure 4A). Given eIF4E, RPS6, rRNA and active ribosomes (via VISTA-R) were located in the same pseudopods, we refer to these structures as translationally active pseudopods (T-PODs). It is important to note that there is no expectation that all of a given factor needs to be restricted to T-PODs to have functional consequences for localized translation there. Indeed, we observed eIF4E, active ribosomes, rRNAs both outside of and within T-PODs consistent with activities in both cytoplasmic and localized translation (Figure 4D-F). In all, we observed Ezrin-positive translation complexes in T-PODs suggestive of a role for Ezrin in translating factors directly involved in cell movement to provision T-PODs for motility.

### Translation in T-PODs is Ezrin- and eIF4E-dependent

We next determined the impact of Ezrin on translation. To test whether Ezrin influenced global protein production, we monitored new protein synthesis using puromycilation assays and studied suspension cells because of the material required. We observed that Ezrin knockdown using siRNA did not influence global protein production when compared to negative control knockdown to luciferase (siLUC) by western blot (Figure 5A left panel). For comparison, eIF4E reduction using MM6 CRISPR-4E versus MM6 CRISPR-CTRL cell lines demonstrated reduced new protein synthesis as measured by puromycin incorporation by ∼50% similar to previous studies using eIF4E antisense oligonucleotides and ^35^S-methionine incorporation^65^ (Figure 5A, right panel). Importantly eIF4E only influences the translation efficiency of a subset of mRNAs, mainly those containing long/structured 5’UTRs, which often correspond to mRNAs that encode oncogenic proteins^30, 66^. Indeed, many transcripts such as β-Actin, Tubulin and GAPDH are not sensitive to modulation of eIF4E levels^22^. In all, genetic reduction in Ezrin levels did not have global impacts on new protein production consistent with its restricted localization, suggesting that it impacts a smaller repertoire of mRNAs targets than eIF4E, likely with specialized functions in motility.

**Figure 5.**
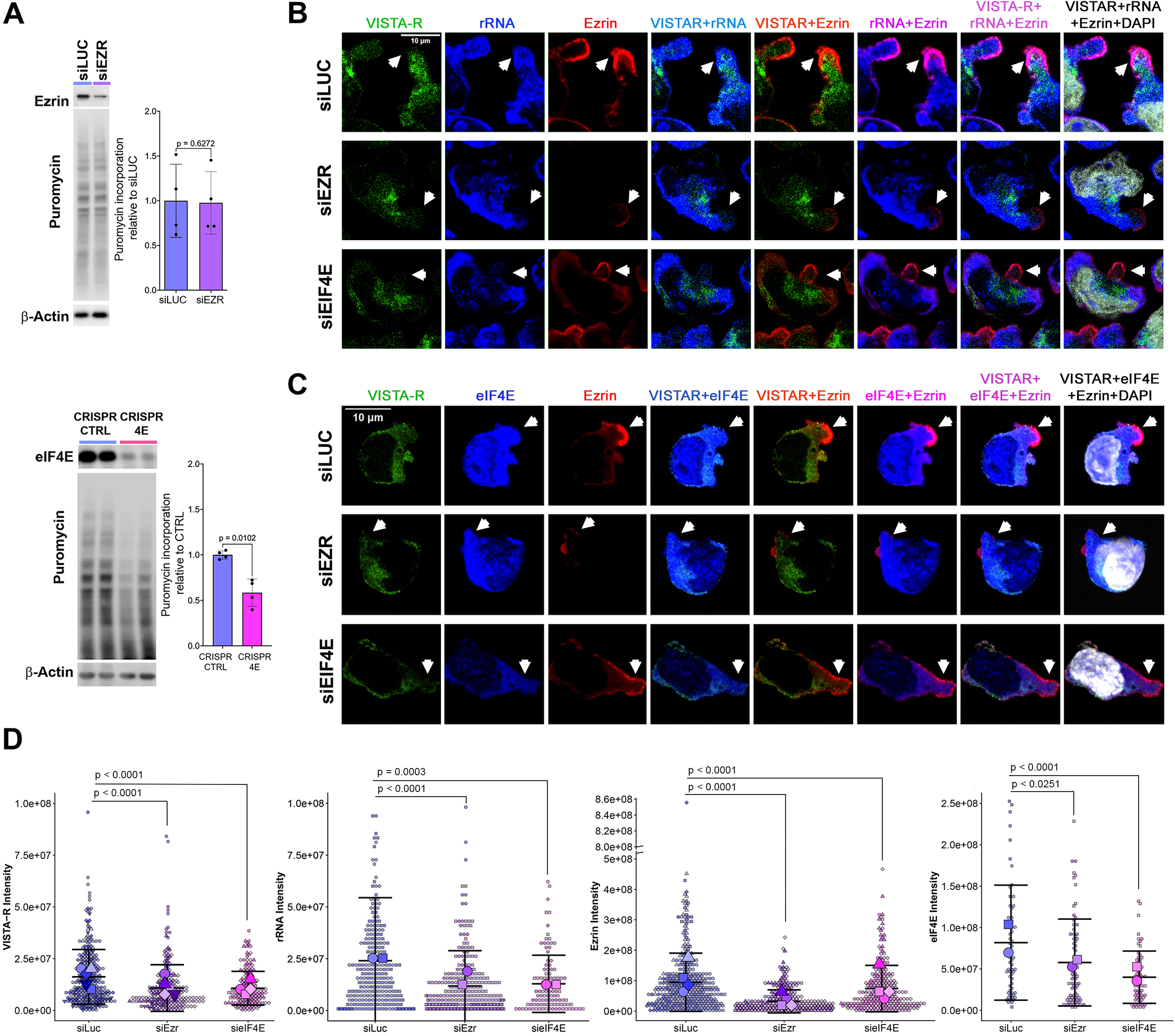
Ezrin influences localized translation in T-PODs but not global translation. **A.** Western blots of total cell lysates for MM6 cells in suspension treated with puromycin and analyzed by WB with anti-puromycin antibody (to assess global protein synthesis) reveal that Ezrin does not influence global translation: RNAi knockdown to Ezrin (siEZR) did not show difference compared to luciferase (siLUC) control (left panel). In contrast, CRISPR 4E reduces global translation signal by about 50% relative to CRISPR-CTRL, consistent with previous studies^30, 65^. Each symbol represents a biological replicate, bar showing mean, standard deviation, p-value (two tailed paired t test). **B.** Confocal micrographs for VISTA-R and the indicated proteins for MM6 cells upon genetic knockdowns siEIF4E, siEZR or siLUC control. Arrows indicate T-PODs. Scale bar = 10µm. Western blots confirmed knockdown (Supplementary Figure 6C) **C.** As in B but monitoring eIF4E in T-PODs as a function of Ezrin and eIF4E knockdown. Ezrin is also reduced by eIF4E knockdown in invaded cells (Supplementary Figure 6C). Scale bar = 10µm. **D.** Imaris-quantified intensities of VISTA-R, rRNA, Ezrin, and eIF4E within T-PODs under the indicated conditions. Data are presented as superplots. Each small symbol represents an individual cell; different shapes represent independent experiments, with larger symbols indicating the mean for each individual experiment. Horizontal lines indicate the mean ± standard error across all independent experiments. P-values were calculated using Welch’s t-test.

Next, we determined whether Ezrin influenced localized translation in T-PODs using VISTA-R in invaded cells. To do this, siRNA knockdown of Ezrin and eIF4E were carried out and compared to the negative control siLUC with knockdowns validated by western blot (Supplemental Figure 6C). Confocal microscopy was used to monitor localization in T-PODs for active ribosomes (VISTA-R), rRNA (Y10b), Ezrin and DAPI (Figure 5B) and VISTA-R, eIF4E, Ezrin, CD44 and DAPI (Figure 5C). Arrows indicate example T-PODs. In siLUC controls, we observed a staining pattern similar to that of parental invaded MM6 cells (Figure 5B-C, top row; Figure 4D-F), with Ezrin, eIF4E, rRNA and VISTA-R localized to the same pseudopods (Figure 5B-C, top row). Thus, siLUC treatment did not disrupt the localization of these factors or the translation activity in the T-PODs relative to invaded MM6 cells. By contrast, siEIF4E and siEZR treatment led to a reduction of active ribosomes, Ezrin, eIF4E and rRNAs in T-PODs when compared to siLUC (Figure 5B,C).

To quantify the signal intensity in T-PODs for each channel, we used Imaris software and with the following steps: 1. A surface corresponding to the nucleus was generated using the DAPI signal, 2. The MATLAB XTension *Distance Transformation* was applied to the DAPI-defined surface to generate a new surface capturing channel signals on and within the cellular membrane of each cell (T-PODs remained outside of this surface), 3. All protein signals within the Distance Transformation surface were removed by masking, 4. A new surface corresponding to T-PODs was generated using the newly created channel following masking of the Distance Transformation surface, 5. Signal intensities for each channel were measured within isolated T-POD surface volumes (Supplemental Figure 6D), and presented as calculated intensities are per T-POD. Using this strategy, we observed that eIF4E knockdown significantly reduced the VISTA-R signal and rRNA intensities by 1.5- and 1.9-fold per T-POD relative to siLUC controls, respectively (Figure 5D). eIF4E knockdown also reduced Ezrin levels per T-POD (Figure 5D) consistent with eIF4E reducing Ezrin protein levels in total cell lysates (Figure 1B, 3A, Supplemental Figure 5A, Supplemental Figure 6C). Interestingly, Ezrin knockdown reduced the signal per T-POD from active ribosomes (VISTA-R) 2-fold and rRNA 2.4-fold relative to siLUC controls (Figure 5B-D). For positive controls, signals were reduced for eIF4E (2-fold) and Ezrin (2.3-fold) per T-POD upon their respective knockdowns (Figure 5D). Importantly, although rRNA levels were reduced per T-PODs upon Ezrin or eIF4E knockdown (Figure 5B-D), their overall levels in cells were not altered as measured by RT-qPCR (Supplemental Figure 6E). Given the following findings: (1) Ezrin depletion led to decreased levels of rRNA, active ribosomes and eIF4E per T-POD (Figure 5B-D), (2) Ezrin did not impact global translation, or total levels of eIF4E or rRNA (Figure 3A, Figure 6B, Supplemental Figure 6E), (3) Ezrin binds the transmembrane protein CD44 (Figure 3D)^67^, and (4) Ezrin is enriched in T-PODs (Figures 3M, 4D-F, 5B-D); we concluded that the role of Ezrin is to anchor the active ribosomes to T-PODs. Given eIF4E and Ezrin directly interact in vitro, we postulate that this anchoring is direct but may require additional stabilizing factors which can be investigated in future. Moreover, we demonstrate that the presence of active ribosomes in T-PODs is dependent on both Ezrin and eIF4E.

**Figure 6.**
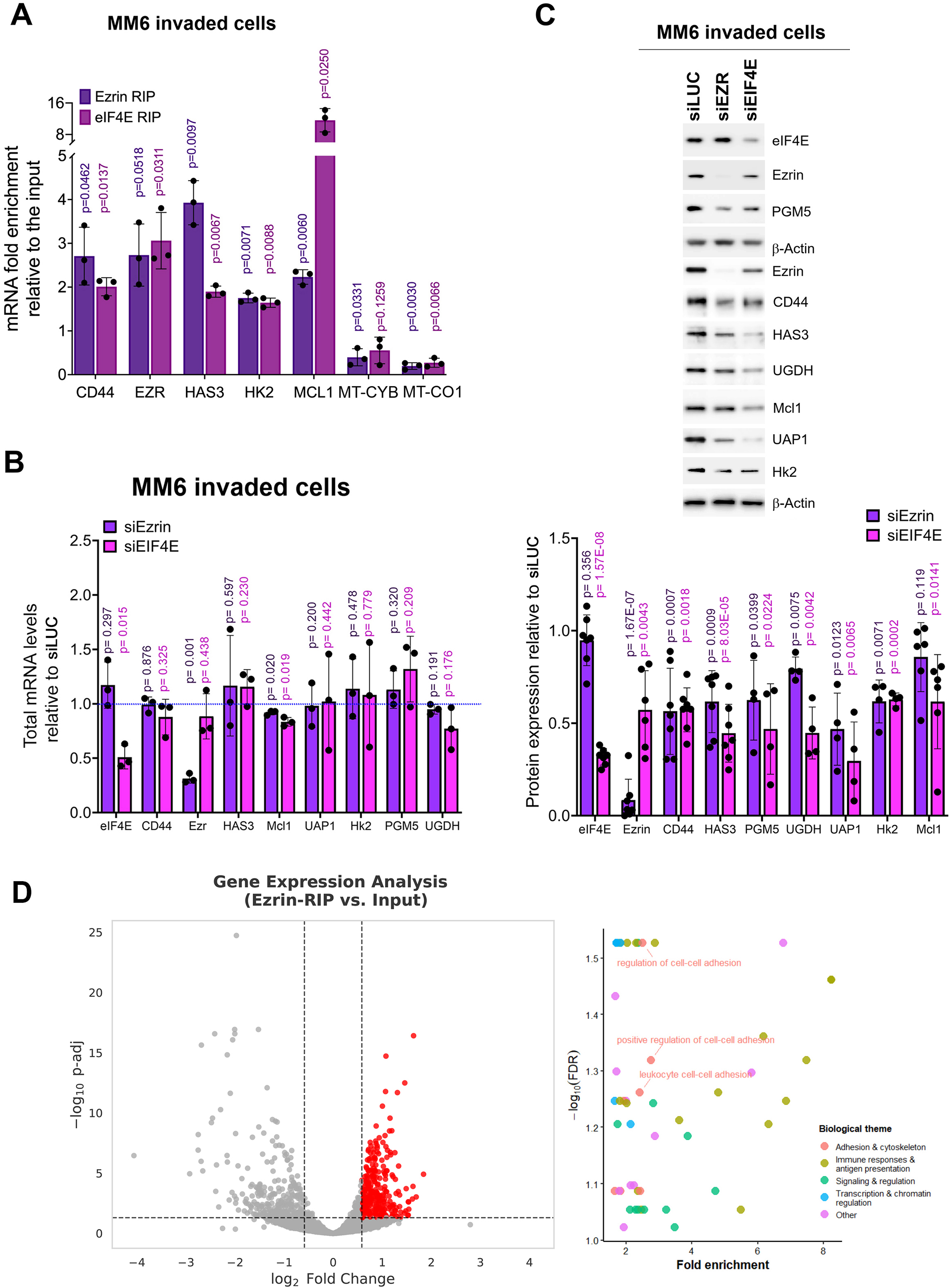
Ezrin physically associates with RNAs encoding motility factors and influences their protein production. **A.** Ezrin and eIF4E RIPs from cytoplasmic fractions from invaded MM6 cells compared to input. Each symbol represents a biological replicate. Means, standard deviations, p-values (Welch’s t test) are shown. Mitochondrial encoded mRNAs were used as negative controls (MT-CYB and MT-CO1). **B.** Total mRNA levels detected by RT-qPCR in invaded MM6 cells as a function of genetic knockdown using siRNA to *EIF4E* or *EZR* as compared to siLUC treated cells. Each symbol represents a biological replicate performed independently. Bars represent the mean, shown with standard deviations and p-values (Welch’s t test). **C.** Top: Representative western blots showing the expression for the indicated proteins using total cell lysates of invaded MM6 cells with knockdown of Ezrin (siEZR) or eIF4E (siEIF4E) and siLUC control. Bottom: Quantitation of indicated proteins analyzed by western blots: each symbol represents a biological replicate, bar shows the mean, standard deviations and p-values (multiple unpaired t tests). **D.** Left: Volcano plot showing transcripts significantly enriched in Ezrin RIP-seq (fold change >1.5 and adjusted *p* value <0.05). Right: Gene Ontology analysis of transcripts significantly enriched in Ezrin RIP-seq.

### Ezrin physically associated with transcripts encoding motility factors and impacted their protein production

While VISTA-R enabled us to observe active ribosomes in T-PODs in fixed cells, identification of mRNAs that underwent translation in T-POD would be challenging. The following issues to identify transcripts that undergo Ezrin-dependent translation needed to be considered in order to develop fruitful solutions: (1) isolation of protrusions after treatment with labeled amino acids and followed by protein analysis would not reliably identify newly synthesized proteins generated in T-PODs given rapid diffusion of protein products from ribosomes (in seconds)^62^ and diffusion between T-PODs and bulk cytosol which would confound assignment of the origin of translation, (2) measurement of bulk translational efficiency using polysomes or ribosome foot-printing methods would measure translation efficiency over the entire cell whereby T-POD translation will influence translation of a much smaller pool (but physiologically important) of mRNAs which would not be detected over the bulk signal, and (3) in invaded cells or in isolated protrusions there is not sufficient material to measure translation efficiency even using low-input polysome analysis methods. Thus, we opted, as a first step, to identify RNA targets of Ezrin-dependent translation which should allow us to identify T-POD translation targets by leveraging the enrichment of Ezrin in these structures in invaded cells (Figure 4D-F) and the physical association between Ezrin, eIF4E and the translation machinery (Figure 3).

With this in mind, we fixed invaded cells in the lower Boyden chamber, collected these, isolated the cytoplasmic fraction which then underwent immunoprecipitations with Ezrin or eIF4E antibodies or the negative control IgG (Figure 6A). Shared targets would provide strong evidence that these are indeed translation targets of Ezrin. Cytoplasmic fractions were used to avoid isolating eIF4E-dependent mRNA export targets from the nucleus; the quality of these fractions is shown (Supplemental Figure 5E). It was only possible to isolate ∼100 µg of invaded cell material per six 6-well chambers which then underwent fractionation and immunoprecipitation. Due to these limiting amounts, not every mRNA was detected in the cytoplasmic input. We also note that in some cases no signal was detected at all in the IgG negative control by RTqPCR, which confirms no background binding, but obviously could not be plotted. Given its role in motility and the dependence we observed above between Ezrin and Ezrin-CD44-HA pathway we monitored transcripts encoding these factors. Ezrin immunoprecipitated *CD44* and *Ezrin* transcripts and mRNAs encoding factors involved in HA biosynthesis *HAS3*, and *HK2* by 2–4-fold relative to input (Figure 6A). We reasoned that Ezrin RIP targets that were translation targets would also physically associate with eIF4E. Consistent with this, eIF4E RIPs with these same targets by ∼2-fold relative to input (Figure 6A). As negative controls, we examined eIF4E and Ezrin RIPs of mitochondrially encoded transcripts cytochrome B (*MT-CYB*) and cytochrome oxidase 1 (*MT-CO1*), consistently neither was enriched in eIF4E nor Ezrin RIPs. Interestingly, Mcl1 was much more enriched in eIF4E RIPs (12-fold relative to input), consistent with eIF4E’s presence in both the cytoplasm (bulk translation) and T-PODs compared to Ezrin RIPs (2-fold versus input), consistent with its enrichment in T-PODs but not in the cytoplasm. We note that we cannot rule out the relative avidity of the eIF4E and Ezrin antibodies as contributors to this observation. We ruled out effects of Ezrin or eIF4E on mRNA stability and/or transcription of these target transcripts since there were no changes to target mRNA levels in siEZR or siEIF4E relative to siLUC conditions examining mRNAs isolated from total cell lysates derived from invaded cells (Figure 6B). Moreover, these knockdowns did not impact the total levels of 18S, 28S or 5S rRNAs (Supplemental Figure 6E) suggesting that the number of ribosomes was not influenced. Thus, Ezrin and eIF4E did not influence the steady-state stability or transcription of these transcripts; and moreover, Ezrin which is substantially enriched in T-PODs is physically associated with these mRNAs. In all, these findings position Ezrin as a positive modulator of the translation of factors in the Ezrin-CD44-HA axis through physical association of Ezrin with eIF4E and through this interaction with target mRNAs.

Next, we sought to determine if Ezrin influenced protein production of these targets (Figure 6C). Here, invaded cells were collected and total cell lysates prepared. By western blot, we observed that Ezrin knockdown was about 10-fold and Ezrin and eIF4E knockdown was 3-fold compared to siLUC. siEZR decreased protein levels by ∼40-60% for CD44, HAS3, HK2, PGM5, and UAP1 proteins and around ∼25% for UGDH compared to siLUC (Figure 6C). MCL1 trended to ∼20% reduction upon siEZR relative to siLUC controls without reaching statistical significance.

Importantly, reduction in Ezrin did not affect eIF4E protein levels (0.95±0.15 siEzr compared to siLUC, p-value=0.356) consistent with the same observation for cells in suspension (Figure 6C, Figure 3A, Supplemental Figure 5A). We anticipated greater effects for eIF4E than Ezrin on target protein production despite the 3-fold reduction in eIF4E relative to the 10-fold reduction in Ezrin. This is because: 1. eIF4E plays a direct role in the nuclear export of these target mRNAs thereby increasing their availability to the ribosome^31^, 2. eIF4E is found throughout the cytoplasm and thus can influence translation outside of T-PODs, and 3. eIF4E reduction decreased Ezrin protein levels by ∼50%. Indeed, the 3-fold knockdown of eIF4E reduced protein levels for CD44, HAS3, HK2, UAP1, PGM5, UGDH and MCL1 by ∼40-70% in invaded cells relative to siLUC controls while the 10-fold reduction of Ezrin was similar but not as marked ranging from 30-60% (Figure 6C). Similar findings were observed with CRISPR mediated knockdown of eIF4E in invaded cells and in cells grown in suspension (Figure 1B, Supplemental Figure 1D, Supplemental Figure 6F), demonstrating the very robust impact of eIF4E on these pathways. In all, impact of Ezrin and eIF4E were highly similar in terms of shared targets in the Ezrin-CD44-HA axis but with 3-fold eIF4E knockdown consistently causing greater effects than the 10-fold reduction of Ezrin as expected. Consistent with a model whereby Ezrin binds eIF4E-m^7^G capped mRNAs complexes through eIF4E (Figure 3K and L), we observe complete overlap between the Ezrin and eIF4E mRNA clients identified and impacts on corresponding protein production. Taken together with our findings that Ezrin directly interacted with eIF4E, physically associated with rRNAs, ribosomal proteins, translation initiation factors, and that Ezrin colocalizes with eIF4E and active ribosomes in T-PODs, our studies revealed an unexpected role for Ezrin in modulation of translation of specific mRNAs encoding motility factors in T-PODs.

To identify other mRNAs that co-exist in Ezrin complexes, we performed Ezrin RIP-Seq on cytoplasmic fractions from MM6 and MM6 CRISPR-CTRL cells (as biological replicates) in suspension. Similar experiments in invaded cells revealed insufficient material for this type of study. Differential enrichment analysis revealed 335 transcripts with a fold change >1.5 and an adjusted *p* value <0.05. Consistent with our findings that Ezrin contributes to the localized translation of proteins required for cell invasion, gene ontology analysis revealed that Ezrin RIPs included mRNAs encoding proteins involved in cytoskeletal remodeling and protrusion formation (e.g., EZR, PLEC, MYH9), leukocyte adhesion and migration (e.g., CCDC88B, ITGB2, ITGAL), and extracellular matrix remodeling (e.g., MMP14, PLAUR), leukocyte cell–cell adhesion, cell growth, innate immune regulation, antigen processing and presentation, as well as RNA splicing and mRNA processing and chromatin remodeling (Figure 6D, Supplementary Table 1). In this way, Ezrin, in cooperation with eIF4E, is positioned to regulate the translation of specific mRNAs to T-PODs. This would allow translation close to the site of function thereby provisioning T-PODs with the proteins required for the invasive phenotype.

## Discussion

Our studies establish a direct physical, spatial and functional link between the motility and translation machineries, centered on a previously unrecognized role for Ezrin. We show that eIF4E drives AML cell motility and disease progression across genetic, pharmacological, and *in vivo* models; that high-eIF4E AML cells depend on Ezrin for this motility while eIF4E-depleted cells do not; and that Ezrin achieves this by binding eIF4E-m^7^G-capped mRNA complexes through its FERM domain, anchoring active ribosomes in T-PODs to sustain on-demand, localized translation of motility factors to produce CD44 and the HA biosynthetic enzymes. Several lines of evidence converge on this model. Ezrin and eIF4E co-immunoprecipitated independently of RNA, and purified Ezrin-FERM directly binds purified eIF4E, indicating a protein-protein rather than RNA-bridged interaction. Cap-affinity chromatography demonstrated that Ezrin is captured on beads by eIF4E in high-eIF4E AML cells arguing that Ezrin engages mRNA indirectly through eIF4E rather than binding transcripts on its own. Importantly, the biochemical data establish physical association of Ezrin with mRNA and ribosomes while the confocal data provided the spatial evidence that these interactions are occurring in T-PODs with the majority of enrichment of Ezrin with the translation machinery at the tips of the T-PODs in motile cells. Because the Ezrin-FERM domain is structurally conserved across the ERM family, the other members (radixin, moesin) may share this capacity which will be explored in future. In all, Ezrin through its direct interaction with eIF4E associated with active ribosomes, ribosomal proteins, rRNAs and mRNAs encoding itself and members of the HA-CD44-Ezrin pathway and influences their translation.

These studies required resolving a methodological problem: puromycin- and label-based assays report where newly made protein ends up, not where translation occurred, because newly synthesized proteins diffuse from ribosomes within seconds. VISTA-R sidesteps this by fixing ribosomes *in situ* before labelling, marking the ribosome itself rather than its product. This is what allowed us to distinguish two competing possibilities, that T-PODs are sites of active translation versus storage depots for translation factors, in favor of the former. Moreover, VISTA-R can be used in fixed tissues where diffusion-based and large-tag fusion methods to visualize in situ translation cannot be applied.

The model that Ezrin anchors eIF4E and active ribosomes in T-PODs predicts, and our knockdown data confirm, a dissociation between global and local translation effects: Ezrin loss did not reduce bulk protein synthesis or total rRNA levels but did reduce active ribosome, rRNA, and eIF4E localization specifically within T-PODs, and reduced protein production of Ezrin-CD44-HA pathway proteins without changing their mRNA transcription or stability. This pattern, normal transcript levels alongside reduced localization of active ribosomes and reduced protein output, is consistent with a role for Ezrin in concentrating existing translational capacity at the T-POD rather than in regulating transcript abundance or global translation output. It also explains why eIF4E depletion, which affects both nuclear export of these mRNAs and their translation, produces larger effects than Ezrin depletion, which acts only at the local translation step; the two are sequential rather than competing mechanisms. That the Ezrin-CD44-HA axis is itself the target of this translation program is notable, suggesting that Ezrin can feed-forward production of motility factors in its own pathway to sustain movement. Moreover, our findings demonstrate that genetically targeting Ezrin shuts down this translation program. Additionally, Ezrin RIP-seq revealed enrichment of a distinct subset of cytoplasmic transcripts encoding proteins involved in cytoskeletal remodeling, adhesion, migration, and extracellular matrix interactions, potentially extending Ezrin’s role in regulation to factors beyond its own pathway. This finding is consistent with the requirement for coordinated production of proteins that enable dynamic reorganization of the cytoskeleton, interaction with the extracellular environment, and formation of invasive protrusions required for an invasive phenotype.

These findings extend our earlier work showing that eIF4E drives nuclear export of *CD44* and HA-pathway mRNAs, increasing their cytoplasmic availability. The present findings identify a second, mechanistically distinct step acting on the same transcripts: once exported, these mRNAs undergo Ezrin- and eIF4E-dependent translation specifically at T-PODs. eIF4E therefore acts on this pathway at two sequential points, nuclear export and local translation, consistent with eIF4E depletion producing a more pronounced phenotype than Ezrin depletion. Ezrin’s reported interactions with PABP1 and DDX3 in stress-granule contexts raise the possibility of a broader role in RNA metabolism beyond the T-POD mechanism described here; we did not observe stress granules in our system, which may reflect the absence of the stress conditions under which they were previously reported rather than their absence from this pathway^68^.

Like many other groups that have faced these same challenges studying localized translation^69, 70^, which in this case is further exacerbated here by the low number of invaded cells, we directly measured production of the target proteins. However, we extended this by monitoring protein targets whereby their corresponding mRNAs were bound to Ezrin and eIF4E, demonstrating a clear link between target binding and corresponding protein production. Indeed, even current low input polysome analyses are not feasible with the material available. In future, the development of more sensitive detection methods for translation efficiency coupled with ones that can be conducted *in situ* will make direct assessment of ribosomes per transcript in pseudopods feasible.

These are the first studies in mouse models to show that eIF4E played a role in leukemogenesis. Reduced eIF4E levels impaired bone marrow engraftment and improved survival across three independent cell-line-derived xenograft models and a patient-derived xenograft model treated with the eIF4E inhibitor ribavirin. eIF4E-dependent motility in AML cells was Ezrin-dependent. The interactions with CD44 and HA, suggests that these routes could be combined with eIF4E inhibition in future. These findings are consistent with previous observations that elevated eIF4E alone predicted poor outcomes in AML patients (Leucegene.ca)^15^ ^10^. These studies are the first to make a direct link between eIF4E, the motility machinery and localized translation. Previous studies support these findings single staining experiments showed that eIF4E was at the migrating edge of cells^71^. The authors concluded that either there is active translation at these structures or that there is a free pool of eIF4E or a pool of repressed eIF4E in the form of the eIF4E-4EBP1 complex^71^. They conclude that they “were unable to demonstrate preferential synthesis of proteins or active signs of translation”. In another study, newly synthesized proteins were monitored via puromycilation, however there was the rapid diffusion of these signals from ribosomal proteins used to mark translation sites, consistent with the limitations of this method for identifying translation sites^72^. This limitation is overcome with the VISTA-R method since it marks active ribosomes not newly synthesized proteins. In all, we provide novel insights into cell movement demonstrating that AML cell motility was driven at the post-transcriptional level by eIF4E, providing an important addition to the overarching mechanisms underpinning these processes beyond transcription and stability.

In conclusion, we identify a function for Ezrin in eIF4E-dependent localized translation, show that this function is required for eIF4E-dependent AML motility, and develop VISTA-R as a tool that made this localized translation visible and also is usable in settings inaccessible for conventional in situ translation assays such as fixed patient tissues. Together these findings establish an on-demand translation system that provisions migrating AML cells with motility factors at their site of action, and position this system, alongside eIF4E’s established role in mRNA export, as a therapeutic target in AML.

## Materials and Methods

### Cell lines and primary samples

HS-5 cell line was obtained from the American Type Culture Collection (ATCC). MM6, THP-1, and NOMO-1 cell lines from Leibniz Institute DSMZ-German collection of microorganisms and cell cultures. All cell lines were authenticated and routinely tested for mycoplasma (PlasmoTest™, mycoplasma detection kit, rep-pt1, InvivoGen). HS-5 cells were cultured in Dulbecco’s Modified Eagle’s Medium (DMEM, cat# 11995-065, Gibco) with 10% fetal bovine serum (FBS, cat# 100-106, GeminiBio) and 1% penicillin-streptomycin solution (pen/strep, cat# 30-002, Corning); MM6, THP-1 and NOMO-1 cells were cultured in Roswell Park Memorial Institute medium (RPMI 1640, cat# 11875-093, Gibco) supplemented with 10% FBS and 1% pen/strep. MM6 cells were also supplemented with non-essential amino acids (cat# 11140-050, Gibco), 1 mM sodium pyruvate (cat# 11360-070, Gibco) and 10 µg/ml human insulin (cat# I9278, Sigma-Aldrich).

For eIF4E or Ezrin knockdowns, MM6 cells were transfected with corresponding siRNA duplexes using SF Cell Line 4D X Kit (Lonza) following program Jurkat on the Amaxa® Nucleofector® Technology (Amaxa Biosystems, Berlin, Germany). All siRNA were from IDT Technologies. siRNA duplex_Luciferase sense (CACGUACGCGGAAUACUUCGAAATG) and antisense (CAUUUCGAAGUAUUCCGCGUACGUGUU); siRNA duplex _Ezrin sense (GGAAUCAACUAUUUCGAG) and antisense (UUUUUAUCUCGAAAUAGU); siRNA duplex eIF4E: sense (CUGCGUCAAGCAAUCGAGAUUUGGG) and antisense (CCCAAAUCUCGAUUGCUUGACGCAGUC).

MM6 and THP-1 CRISPR clones with eIF4E disruption were generated using Lenticrisprv2 plasmid purchased from Addgene^73, 74^. Lenticrisprv2 plasmids with sgRNA inserts (eIF4E: TTAAACATCCCCTACAGAAC, and Azami-Green protein as the control: GGCCACAACTTCGTGATCGA) were kindly provided by the lab of Dr. Mike Tyers. MM6 cells (ATCC) were transduced using lentiviral particles containing these plasmids and produced in 293T cells and selected with puromycin at 1µg/ml. Clones were screened for eIF4E expression by western blot. Knockdown in clones with reduced protein expression was confirmed by Sanger sequencing.

NOMO-1 overexpressing cell lines were produced by retroviral transduction. Briefly, eIF4E wild-type retroviral vectors along with pCL-Ampho packaging vector were transiently transfected into 293T cells (ATCC), and retroviral supernatants were used to infect NOMO-1 cells. GFP^+^ cells were isolated using a cell sorter (FACSAria; BD Biosciences).

There were three sources for deidentified primary human specimens all of which were obtained with written informed consent according to the Declaration of Helsinki and IRB/Ethics Committee institutional approvals: Weill Cornell Medicine, Banque de cellules leucémiques du Québec (BCLQ), supported by the Cancer Research Network of the Fonds de Recherche du Québec - Santé (FRQS), or from AML patient screen samples prior to enrollment in our ribavirin combination therapy clinical trial with Health Canada approval^51^. Blasts were isolated using flow cytometry as described^12^. Specimens from healthy donors were obtained from ATCC and Lonza.

### Plasmids, antibodies and reagents

pcDNA-2Flag-eIF4E wild-type and vector as well as MSCV-pgk-GFP-eIF4E and control constructs were previously described^11, 26, 39, 75^.

Antibodies for immunoblotting: mouse monoclonal anti-eIF4E (cat# 610270, BD Biosciences), mouse monoclonal anti-β-Actin (cat# A5441, Sigma Aldrich), rabbit polyclonal anti-Mcl-1 (S-19) (cat# sc-819, Santa Cruz), mouse monoclonal anti-HSP90α/β (F-8) (cat# sc-13119, Santa Cruz), rabbit polyclonal anti-Myc (cat# ab32072, Abcam), Mouse monoclonal anti-CD44 antibody (cat# 156–3 C11, Cell Signaling Technology), rabbit polyclonal anti-CD44 (cat# A12410, Abclonal), rabbit polyclonal anti-HAS3 antibody (cat# ab154104, Abcam), rabbit polyclonal anti-phosphoglucomutase 5 (cat# AI14638, Abgent), rabbit polyclonal anti-Lamin A (C-terminal) (cat# L1293, Sigma Aldrich), rabbit polyclonal anti-GAPDH (FL-335) (cat# sc-25778, Santa Cruz), rabbit polyclonal anti-UGDH (cat#AP12613b-EV, Abgent), rabbit monoclonal anti-Hexokinase II (cat# 2867, Cell Signaling Technology), rabbit monoclonal anti-UAP1 antibody (cat# 2716, GenuinBiotech), mouse monoclonal anti-RPL17 (C-8, cat# sc-515904, Santa Cruz), mouse monoclonal anti-RPS16 (D-8, cat# sc-518206; Santa Cruz), mouse monoclonal anti-RPS6 (C-8, cat# sc-74459, Santa Cruz), mouse monoclonal anti-eIF3e (G-7, cat# sc-390413, Santa Cruz), mouse monoclonal anti-eIF3p110 (B-6, cat# sc-74507, Santa Cruz), mouse monoclonal anti-eIF3b (A-7, cat# sc-374156, Santa Cruz), mouse monoclonal anti-eIF4AI/II (H-5, cat# sc-377315, Santa Cruz), mouse monoclonal anti-eIF4G (A-10, cat# sc-133155, Santa Cruz), mouse monoclonal anti-Nopp140 (E-7, cat# sc-374033, Santa Cruz), rabbit monoclonal anti-histone H3 acetyl K27 (cat# ab177178, Abcam), rabbit polyclonal Calreticulin (cat# AW5211, ABCEPTA), rabbit polyclonal anti-Calnexin (cat# ab22595, Abcam), rabbit polyclonal anti-Ezrin antibody (cat# PA5-80603, Invitrogen), mouse monoclonal anti-Ezrin (CPTC-Ezrin-1, cat# AB_2100318, DSHB), mouse monoclonal anti-MEK-1 (H-8, sc-6250, Santa Cruz), and mouse monoclonal anti-cytochrome c (A-8, cat# sc-13156, Santa Cruz). Secondary antibodies: Peroxidase AffiniPure™ Goat Anti-Mouse IgG (cat# 115-035-146, Jackson Immunoresearch), Peroxidase AffiniPure™ Goat Anti-Rabbit IgG (cat# 111-035-144, Jackson Immunoresearch) and mouse monoclonal anti-puromycin (cat# MABE343, Millipore).

For Immunoprecipitations: rabbit polyclonal anti-Ezrin antibody (cat# PA5-80603, Invitrogen), rabbit polyclonal anti-eIF4E (cat# RN001P, MBL international), mouse monoclonal anti-ribosomal RNA (Y10b, cat# AB_2313703, DSHB), Normal rabbit polyclonal IgG (cat#12-370, Millipore), Normal mouse IgG (cat#12-371, Millipore).

For immunostaining: rat monoclonal anti-hum/mouse CD44 (IM7; cat# 14-0441-82, eBioscience), mouse monoclonal anti-ribosomal RNA (Y10b, cat# AB_2313703, DSHB), rabbit polyclonal anti-Ezrin antibody (cat# PA5-80603, Invitrogen), mouse monoclonal anti-eIF4E (cat#610270, BD Biosciences), rabbit anti-eIF4E (cat# RN001P, MBL international).

Secondary antibodies for immunostaining, all from Jackson Immunoresearch: Rhodamine Red-X-conjugated affinity pure Donkey Anti-Mouse IgG (cat# 71529515), Alexa Fluor® 647 AffiniPure Donkey Anti-Rat IgG (H+L) (cat# 712-605-153), Alexa Fluor® 488 AffiniPure Donkey Anti-Mouse IgG (cat# 715-545-151), Alexa Fluor® 488 AffiniPure Goat Anti-Mouse IgG, Fcγ subclass 2a specific (cat# 115-545-206), Brilliant Violet™ 480-conjugated AffiniPure Donkey Anti-Rabbit IgG (cat# 711-685-152), Rhodamine Red-X-conjugated affinity pure Donkey Anti-Mouse IgG, subclass 1 specific (cat# 115-295-205), Alexa Fluor® 790 AffiniPure® Donkey Anti-Rat IgG (cat# 712-655-153), Alexa Fluor® 647 AffiniPure® Donkey Anti-Rabbit IgG (cat# 711-605-152).

Ribavirin was purchased from Kemprotek (cat# 36791-04-5), puromycin from Santa Cruz (sc-108071B), Homoharringtonine from MedChemExpress (cat# HY-14944). Hyaluronidase from *Streptomyces Hyalurolyticus* was obtained from Millipore (cat# H1136).

### Western blot analysis

Western blots were performed as described previously^31^. Briefly blots were blocked in blocking buffer containing 5% milk in TRIS buffered saline (TBS)–Tween 20. Primary antibodies were diluted in blocking buffer. Quantification of blots was carried out using Fiji (doi:10.1038/nmeth.2019).

### Invasion and adhesion assay

Seventy-two hours prior to the assay, HS-5 cells were seeded into 24 well transwell inserts (5x10^4^ cells per well). 24 hours prior to the assay, MM6 cells were seeded in fresh media at near confluent densities (8x10^5^/mL), in 5-10 mL volume. After growing in fresh media for 24 hours, cells were harvested and washed twice with 1xPBS. Cell pellets were resuspended in 10 mL of 5 μM DiIC12(3) perchlorate (cat# ENZ-52206, Enzo Life Sciences) diluted in RPMI medium without additives (1:10 000 from a 50 mM stock solution) and incubated for 10 minutes at 37 °C. After washing three times with 1X PBS, cells were resuspended in 5 mL of complete RPMI medium, counted, diluted to 3 x10^5^ cells/mL and 0.5 mL of cell suspensions were seeded into the upper chamber of 24-well transwell fluoroblok inserts (with an 8 μm pore size PET membrane, Cat#351152 Corning) in triplicates, and in parallel the same number of cells were plated into 24-well plates at same density and volume as for upper chamber of fluoroblock inserts. Complete growth RPMI medium (0.75 mL) containing 50 ng/mL MCP-1 (279-MC, R&D Systems) was added to the lower chamber. For each experiment plating control wells were seeded, where the same number of cells used for invasion and adhesion assays was seeded in 24-well plates (in technical triplicate for each condition). Thus, we monitored and corrected for errors that could occur in the counting and seeding steps. Cells were incubated for 18-24 hours at 37°C.

For the adhesion assay, HS-5 cells were seeded into 24-well plates 72 hours prior to the assay at 6x10^4^ cells per 24 well. MM6 cells were seeded into a fresh media 24 hours before the experiment as for the invasion assay. After staining with DiIC12(3) perchlorate, cells were seeded into 24-well plates with HS-5 cells at 3 x10^5^ cells/mL and incubated for 1 hour. Cells were gently washed three times with 1XPBS, culture media was added and plates were used for quantification.

For quantification of invasion and adhesion, images of invaded or adhered cells and their plating controls were taken using a 2.5x objective fluorescent microscope set to the Cy3 channel (Leica DM IRB Inverted Fluorescence Microscope®). The captured images were used for quantification using NIH ImageJ® software (https://imagej.nih.gov/ij). Quantification was also done with Cytation 3 plate reader (BioTek).

### Mesenchymal stromal cell spheroids (3D co-culture system)

Spheroids were established as previously reported^41^. Briefly, HS-5 cells were first seeded at 25,000 into 96-well round-bottom plates pre-coated with 1% UltraPure Agarose (cat# 16500, Invitrogen). Twenty-four hours later, when spheroids were formed, 25,000 AML cells were added per well. AML cell lines, that did not express GFP, were pre-stained with CellTrace™ Violet (CTV, cat# C34557, Invitrogen). At 24 hours of co-culture, the spheroid was picked up using a 200 µL-pipette tip, washed with DPBS-5mM EDTA (cat# 15575-038, Invitrogen), and enzymatically disaggregated with 0.25% Trypsin/2.21 mM EDTA (cat# 25-053, Corning) for 10 minutes, followed by mechanical disruption. Single cell suspensions were stained with 7-AminoActinomycin D (7-AAD, cat# A1310, Invitrogen) as viability dye and APC Cy7 anti-human CD45 (hCD45) antibody (cat# 368516, BioLegend).

### AML mouse models

All animal procedures were conducted under protocols approved by the Weill Cornell Medicine Institutional Animal Care and Use Committee (WCM-IACUC). Female, 8-10-week-old, non-obese diabetic/severe combined immunodeficient IL2Rgamma^null^ (NSG) mice were obtained from the Jackson Laboratories (strain #:005557).

For cell line derived xenografts (CDX), MM6 CRISPR cell lines CTRL or 4E clones (5.5 x 10^5^), and THP-1 CRISPR cell lines CTRL or 4E clones (4 x 10^6^) were injected via tail vein into sub-lethally irradiated NSG mice (200 rad). Bone marrow (BM) and spleens were harvested to assess leukemia cell engraftment. Cells were stained with PE-Cy5 anti-mouse CD45 (mCD45) (cat# 103110, BioLegend) and APC Cy7 anti-hCD45 (cat# 368516, BioLegend) antibodies to distinguish human cells from host mouse cells. 4’,6-diamidino-2-phenylindole (DAPI, cat# D1306, Invitrogen) or fixable viability dye eFluor™ 450 (cat# 65-0863-14, Invitrogen) were added to assess viability. Additional monitoring was done using BM aspirates and overall survival was also assessed. Mice exhibiting signs of distress were euthanized in accordance with WCM-IACUC guidelines. We confirmed by western blot that MM6 CRISPR-4E and THP-1 CRISPR-4E cells remained deficient in eIF4E at the endpoint, briefly AML cells (hCD45⁺) were column-purified from mouse BM samples using anti-human CD45 MicroBeads (cat# 130-045-801, Miltenyi Biotec).

AML patient derived xenografts (PDX) models were established based on previous studies^76, 77^. Briefly, primary AML cells (2.5 × 10⁶ cells per mouse) were injected into sublethally irradiated NSG mice (200 rad). Seventeen days post-cell injections, mice were treated via intraperitoneal injection with either 100 mg/kg of Ribavirin (cat# S2504, Selleck Chemicals) or vehicle control (saline) daily for 28 days, followed by a 44-day drug-free period. Treatment cycle was then repeated using 100 mg/kg for 20 days. Peripheral blood was periodically monitored to assess leukemia burden, as described previously^77^. Overall survival was also evaluated.

### Immunofluorescence and laser-scanning confocal microscopy

MM6 cells grown in liquid cultures (suspension) or collected from the bottom chamber of transwell system from the invasion assays (invaded; as shown in Supplemental Figure 6A) were fixed with 3% paraformaldehyde (PFA, cat# 158127, SIGMA) for 15 minutes at room temperature (RT) with gentle shaking (PFA was added directly to the growth media), followed by quenching with 150mM glycine (BP381-5, Fisher) for 5 minutes at RT with gentle shaking (glycine was added directly to cells in media with PFA). Cells were harvested and washed three times in 1X PBS in Eppendorf tubes, using centrifugation, 5 minutes at 1200 rpm. Washed pellets were resuspended in 1X PBS at 0.5-1x10^5^ cells/10μl, and 10μl cell suspensions were spotted on glass slides. After drying, slides were washed once with 1 x PBS, permeabilized with 0.5% Triton X-100 (T-9284, Sigma) for 10 minutes at RT, washed three times with 1X PBS, blocked for 1 h in blocking solution (10% FBS and 0.1% Tween 20 in PBS) and incubated with primary antibodies diluted in blocking solution overnight at 4°C. After washing, cells were incubated with fluorescently labeled secondary antibodies diluted in blocking solution for 1h at RT, washed three times in 1XPBS and mounted in ProLong^TM^ gold antifade media with DAPI (cat# P36935, Invitrogen).

Imaging was carried out using the Zeiss LSM700 with 63x oil objective and numerical aperture of 1.4 (Figures 1A and 1B, Supplemental Figure 1E); Leica TCS SP8 DLS line scan confocal microscope, with 63x oil objective and numerical aperture of 1.4 (Figure 3M, Supplemental Figure 1B, with z-stack step 0.3 µm); the Nikon AXR confocal microscope with 60x oil objective and numerical aperture of 1.42 (Figures 4D and 5B, Supplemental Figure 6B, with z-stack step 0.2 µm); or the Nikon CSU-W1 Dual CAM spinning disc microscope with 60x oil objective and numerical aperture 1.42 (Figures 4E, 4F, 5C, with z-stack step 0.3 µm) or 100x oil objective and numerical aperture 1.45 (Figure 4B, with z-stack step 0.3 µm). Channels were detected separately, with no crosstalk observed. Confocal micrographs represent single sections through the plane of the cell. Confocal micrographs represent single sections through the plane of the cell. Images were obtained from ZEN (Carl Zeiss, Inc.) or FIJI software.

### Visualizing Translation Activity using RiboLace (VISTA-R) staining

MM6 cells were fixed and permeabilized as described in the previous section (see Supplemental Figure 6A). After washing with 1X PBS, cells were then washed twice with RL buffer (20mM HEPES pH=7.4, 10mM MgCl_2_, 137mM NaCl, 2.7mM KCl) and blocked in 2% bovine serum albumin (BSA, cat# BP9703-100, Fisher Scientific) with 50μg/ml streptavidin (cat# 434302, Invitrogen) in RL buffer for 30 minutes at RT. Cells were then washed three times with RL buffer and blocked for 30 minutes at RT with 2% BSA with 2mg/ml Biotin (cat# B1595, Invitrogen) in RL buffer. After washing three times with RL buffer, cells were incubated for 1 hour at RT with 50 µM Biotinylated Puromycin (cat# 2379889-82-2, MedChemExpress) in RL buffer with 2% BSA. After washing three times with RL buffer, cells were incubated with 2μg/ml NeutrAvidin Oregon Green 488 (cat# A6374, Invitrogen) in 2% BSA in RL buffer for 1 hour at RT, washed three times with RL buffer and either mounted in ProLong^TM^ gold antifade media with DAPI, or stained for proteins as indicated in the text. In the case of immunostaining after VISTA-R, cells were blocked for 1 hour in blocking solution (10% FBS and 0.1% Tween 20 in PBS) and incubated with primary antibodies diluted in blocking solution overnight at 4°C. After washing, cells were incubated with fluorescently labeled secondary antibodies (all from Jackson Immunoresearch) diluted in blocking solution for 1 hour at RT, washed three times in 1XPBS and mounted in ProLong^TM^ gold antifade media with DAPI.

### Imaris-Based Quantification of T-PODs

All image processing and quantitative analyses were performed using Imaris v10.2 (Oxford Instruments). The Distance Transformation XTension (MATLAB) was used for nuclear (DAPI) surface expansion. DAPI surfaces were generated using automatic absolute intensity thresholding without the use of creation parameters and filtering. The distance transformation XTension was applied in Outside SurfaceObject mode to compute a distance channel from nuclear boundaries. A distance surface was generated using threshold bounds of 0.00-1.70, capturing cytoplasmic regions from the nuclear envelope to the plasma membrane. Ezrin, VISTA-R, and rRNA (Y10b) surfaces were generated using background subtraction (Local Contrast) with a largest sphere diameter of 10 µm and automatic Absolute intensity thresholding. Object-Object statistics was disabled. Surfaces were filtered to remove background, multicellular aggregates, and non-viable cells. Filtering parameters differed slightly between experimental conditions (CTRL 7 vs. CTRL 13) based on different staining days but followed consistent biological criteria: retention of single, viable cells completely in view of the image file. For CTRL 7, Ezrin surfaces were filtered using a voxel number threshold of > 8,000, area < 950 µm^2^, mean intensity (CD44 channel) > 100, and minimum intensity (Ezrin channel) > 100. VISTA-R and rRNA surfaces were filtered more stringently to account for additional background (voxel number > 1.5×10^4^ for VISTA-R and > 2.7×10^4^ for rRNA; area 150-950 µm^2^), with additional spatial constraints excluding objects within 3-5 µm of image borders in the X and Y dimensions, if necessary. To exclude DAPI-positive non-viable cells, which exhibited abnormal nuclear signals, summed and maximum intensity thresholds were applied to the relevant DAPI channels (Intensity Sum > 1.47×10^7^ and Intensity Max > 1.00×10^-5^). These thresholds were empirically defined from the distribution of nuclear intensities across the dataset and removed cells including atypical nuclear signals. For CTRL 13, Ezrin, VISTA-R and rRNA surfaces were filtered using a minimum voxel threshold of > 8,000, area < 950 µm^2^, mean intensity (CD44 channel) > 100, and minimum intensity (Ezrin channel) > 100, without additional spatial or viability-based intensity constraints, as they were unnecessary. To isolate T-POD associated signal, Ezrin, VISTA-R and rRNA channels were masked using the distance surface by setting voxel intensities inside to 0.00, producing nucleus- and perinuclear-excluded (T-POD) channels (Supplemental Figure 6D). Final surfaces were refined by excluding zero-intensity objects (mean intensity > 0.00001) corresponding to the newly generated T-POD channels. Surface-based measurements were extracted using the Vantage module.

Quantification was performed on Ezrin, VISTA-R, and rRNA surfaces using their corresponding T-POD channels. Identical segmentation and filtering parameters were applied across all image files within the same experimental condition and acquisition time. Intensities per T-POD per channel was calculated using intensity Sum.

### Reverse transcription and quantitative PCR

RNA samples were reversed transcribed using SuperScript VILO (cat# 11754–050, Invitrogen) and Superscript IV cDNA (cat# 11756050, Invitrogen) synthesis kits (for RIP experiments) or MMLV reverse transcription kit using oligo-dT or Random hexamers (cat# 28025013, Invitrogen) for analysis of total RNA levels. qPCR analyses were performed using PowerUp Sybr Green Master Mix (cat# A25777, Applied Biosystems) or Sybr Green qPCR Master Mix (cat# GK10002, GLPBio) in Applied Biosystems Viia7, QuantStudio7 or QuantStudio7 thermal cyclers using the relative standard curve method (Applied Biosystems User Bulletin #2). All the primers are listed in Supplemental Table 2. Cycling conditions: 5 minutes at 95°C followed by 40 cycles of denaturation at 95°C for 5 sec and annealing and extension at 60°C for 30 seconds except for Hk2 and HAS3 primers which were carried at 62°C for invaded cells which had low RNA content.

### Crosslinking and fractionation of cells for IPs and RIPs

For IPs from crosslinked cells, 1-2x10^7^ cells in suspension were fixed with 1% PFA final concentration (directly added to culture media) and incubated with slow shaking 15 min at RT. PFA was quenched with 0.15 M Glycine (final concentration, directly added to cell suspension) and incubated 5 min at RT with slow shaking. For total cell lysates, after washing three times with 1X PBS, cell pellets were resuspended in 1ml NT-2 buffer and divided into two Eppendorf tubes (1.5 ml) and sonicated three times with 5 seconds bursts (30 second pause between each burst) using microtip at 25% power (Sonic Dismembrator Model 500, Fisher, Max Output 400W). For fractionation of crosslinked cells, ∼3x10^6^ fixed cells were resuspended in 0.5 ml NT-2 buffer and sonicated with three rounds of 5 seconds bursts / 30 seconds pause with microtip at power 12. Lysates were centrifuged 3 minutes at 1000xg, cytosolic fraction was transferred into fresh tube, and pellets (nuclei) were pooled and resuspended in 100 µl of NT-2 buffer and sonicated three times 5 seconds burst / 30 seconds pause with microtip at power 20. Nuclear and cytoplasmic lysates were centrifuged for 10 minutes at 10 000xg and supernatants were transferred into fresh tubes. NT-2 buffer: 150 mM NaCl, 50 mM Tris-HCl (pH 7.4), 2.5 mM MgCl_2_, 0.05% Nonidet P-40, supplemented with 1mM DTT, 2x protease inhibitors without EDTA, 200U/ml RNaseOut. Nuclear lysates were centrifuged at 10 000 x g for 10 minutes, and supernatants were transferred into fresh tubes.

### Co-Immunoprecipitation and RIP

Total cell lysates or cytoplasmic fractions (from cells fixed with PFA) adjusted to be no more than 1 mg/ml concentration, were pre-cleared with 50 μL protein G conjugated superparamagnetic beads (Dynabeads Protein G, ThermoFisher Scientific) for 30 min at 4°C. Pre-cleared lysates (1mg total cell lysates or 100-500 µg cytoplasmic lysates) were incubated with 10μg of anti-eIF4E antibody (RN001P, MBL), 8 μg of anti-Ezrin antibody (cat# PA5-80603, Invitrogen), or 10 μg of rabbit IgG (cat# 12-370, Millipore) as a control, and 0.5 mg/ml yeast tRNA (cat# R8508, Sigma Aldrich), overnight at 4°C with rotation. After ON incubation, 50 μl of Dynabeads (cat# 1004D, Invitrogen) were added and incubated for additional 3h at 4°C with rotation. Beads were washed once with NT-2 buffer Supplemented with 1mg/mL heparin (cat# 353003, Sigma-Aldrich) for 5 minutes at 4°C with rotation, and an additional six times with NT-2 buffer with 300 mM NaCl. In case of Y10b IPs, 100 μl of Y10b mouse hybridoma supernatant (DSHB), or 10 μg of mouse IgG (cat# 12-371, Millipore) were bound to 50 μL protein G conjugated superparamagnetic beads in 1 ml of NT2 buffer ON at 4°C, washed three times with NT2 buffer, added to 1 mg of total cell lysates, incubated overnight at 4°C with rotation and washed as described for IPs with rabbit antibodies. After washing, beads were resuspended in 2 X Laemmli buffer (cat# 1610737, Bio-Rad) with β-mercaptoethanol and incubated for 5 minutes at 98°C. Co-immunoprecipitated proteins were resolved on SDS-PAGE and visualized by western blotting. To isolate RNAs from immunoprecipitated reactions, beads were resuspended in elution buffer (100 mM Tris-HCl (pH 6.8), 4% (w/v) SDS, 20% (v/v) glycerol, 12% (v/v) β-mercaptoethanol (444203, Sigma), and incubated for 5 minutes at 98°C. RNAs were isolated using TRIzol reagent (cat# 15596018, Invitrogen) and Direct-zol RNA Micro-prep Kit (cat# R2050,Zymo Research).

### Expression and purification of the GST-FERM

The human Ezrin FERM domain (residues 1–308) was expressed as a GST fusion protein using a pGEX6P-1 plasmid provided by Dr. Sébastien Carréno (Institute for Research in Immunology and Cancer, Canada). *E. coli* BL21(DE3) cells carrying the plasmid were grown in LB medium supplemented with 0.1 mg/mL ampicillin at 37°C. At an OD₆₀₀ of 0.8, expression was induced with 1 mM IPTG overnight at 20°C. Cells were harvested by centrifugation at 6,000 rpm for 20 min at 4°C, and pellets were frozen at −80°C for 20 min before lysis.

Cell pellets were resuspended in cold 1× TBS supplemented with 7 mM β-mercaptoethanol, 2 mM EDTA (pH 8.0), ¼ cOmplete™ Protease Inhibitor Cocktail, 0.008 U/μL DNase, 0.004 mg/mL RNase A, and 0.1% Triton. Cells were sonicated on ice, incubated with 1 mg/mL lysozyme for 15 min, and centrifuged at 20,000 rpm for 30 min at 4°C. Cleared lysates containing GST-Ezrin were incubated with glutathione Sepharose 4B resin (1.5–2.0 mL resin per liter of culture) (cat# 17075605, GE Healthcare) for 2 hours at 4°C. The resin was washed four times with 20 mL cold wash buffer (1× PBS supplemented with 500mM NaCl, 7mM β-mercaptoethanol), followed by 20 mL low-salt buffer (50 mM sodium phosphate (pH 7.2), 50 mM NaCl, 1 mM DTT). GST-FERM was eluted from the resin five times with 1 mL aliquots of elution buffer (low-salt buffer supplemented with 50 mM reduced glutathione) for further purification by two sequential chromatography steps. GST-FERM was first subjected to anion-exchange chromatography using a Mono Q column with a gradient between low- and high-salt buffers (50 mM sodium phosphate (pH 7.2), 1 M NaCl, 1 mM DTT). Fractions corresponding to the GST-FERM peak were analyzed by SDS-PAGE, and fractions exhibiting the highest purity were pooled and further purified by size-exclusion chromatography using a Sephadex 200 column.

### Expression and purification of recombinant eIF4E

Human recombinant eIF4E was expressed and purified as previously described^27^. Briefly, *Escherichia coli* BL21(DE3) cells carrying the pET-28a eIF4E plasmid with a TEV protease cleavage site were grown at 37°C and induced with 0.5 mM IPTG at OD₆₀₀ of 0.7 overnight at 30°C. Cells were harvested by centrifugation and resuspended in PBS supplemented with 300 mM NaCl, 10 mM β-mercaptoethanol, 0.5% IGEPAL, 1 mg/mL lysozyme, ¼ cOmplete™ Protease Inhibitor Cocktail, and 10 mM imidazole. Cells were lysed by sonication on ice, and lysates were centrifuged for 30 min at 20,000 rpm at 4°C. Supernatants were applied to Ni-NTA beads (approximately 1 mL resin per liter of culture), (cat# 88221, Thermo Scientific), washed extensively with lysis buffer containing 20 mM imidazole, and eluted with lysis buffer containing 500 mM imidazole. TEV protease was added to the eluate, and samples were dialyzed against PBS supplemented with 1 mM DTT using three dialysis steps of at least 4 h each, including an overnight incubation. TEV cleavage left two additional residues (Gly-Ser) at the N terminus of eIF4E. Proteins were further purified by size-exclusion chromatography using a Superdex 200 pg column in 50 mM sodium phosphate (pH 7.2), 100 mM NaCl, and 1 mM DTT, yielding purity >95%.

### GST pull-downs

Glutathione Sepharose 4B beads (cat# 17075605, GE Healthcare) were equilibrated by washing twice with 1 mL PBS and once with 1 mL BP-1 buffer. GST or GST-Ezrin-FERM proteins were immobilized at 10 pmol protein per 1 μL beads in 1 mL BP-1 buffer for at least 30 min or overnight at 4°C with rotation. BP-1 buffer: 50 mM sodium phosphate (pH 7.2), 100 mM NaCl, 1 mM DTT, supplemented with 2x protease inhibitors without EDTA. Beads were washed three times with BP-1 buffer.

For each binding reaction, 10 μL of bead slurry containing 100 pmol of GST fusion protein was incubated with the eIF4E in 500 μL HEPES buffer (20 mM HEPES (pH 7.4), 110 mM potassium acetate, 2 mM magnesium acetate, 1 mM EGTA, 20 mM DTT, 0.3% IGEPAL, 2x protease inhibitors without EDTA l) for 2 h at 4°C with rotation. Following binding, beads were washed five times with HEPES buffer. All bead washes were performed with centrifugation at 500 × g for 2 min at 4°C. Washed beads were adjusted to 20 μL and mixed with 30 μL Laemmli sample buffer. Bound proteins were eluted with hot Laemmli buffer. Samples were briefly centrifuged at 16,000 × g at RT, and supernatants were collected, resolved by SDS-PAGE, and visualized by western blotting.

### m7GTP Pulldown Assays

20μL of immobilized γ-Aminophenyl-m7GTP (C10-spacer)-agarose beads (Jena Bioscience) were equilibrated with buffer B containing 0.3M NaCl, 0.1M sodium phosphate (pH 7.5), 10μM protease-free BSA and 0.1% IGEPAL. The beads were spun at 0.5xg for 2 mins at room temperature in between each wash. Negative controls of beads were incubated with GST-tagged FERM domain of Ezrin, or GST, in buffer B, meanwhile experimental conditions of beads were initially incubated with eIF4E, in buffer B at room temperature for 30 minutes and rotated at 16.6rpm. Samples were then centrifuged at 0.5xg for 2 mins at room temperature. The samples were removed, and the beads were washed thrice with buffer B, centrifuging between each wash to discard unbound protein. Respective beads were then incubated with m7GTP, or Ezrin FERM-GST, or GST, or GTP, in buffer B for 30 mins at room temperature while rotating at 16.6rpm. The samples were then washed in buffer B with centrifugation at room temperature for 2 mins, at 0.5xg, with subsequent removal of the unbound protein. Respective beads were then incubated with m7GTP, or GTP, in buffer B for 30 mins at room temperature while rotating at 16.6rpm. The samples were then washed in buffer B with centrifugation at room temperature for 2 mins, at 0.5xg, with subsequent removal of the unbound protein. Proteins that remained bound to the beads were eluted by boiling in 2X Laemmli buffer and visualized through Western blotting.

### Statistical analyses

Graphs and statistics were generated using GraphPad Prism Version 10.6.1. Biological replicates and p-values are provided in each figure and the statistical test applied is named in the legend. Superplots were generated using R version 4.4.2.

### RIP sequencing and Gene Ontology analysis

Ezrin RIPs and RNA isolation were performed as described above. Libraries from Ezrin immunoprecipitates and corresponding input RNAs were prepared using KAPA RNA EvoPrep Kit (cat# 10640561702, Roche) and sequenced on an Illumina NovaSeq platform. Reads corresponding to ribosomal RNA (rRNA) were removed by alignment using Bowtie2^78^.

Remaining reads were aligned to the human reference genome GRCh38 using STAR^79^, and uniquely aligned reads were counted using htseq-count with GENCODE v42 gene annotations^80^. Enrichment of RNAs in Ezrin RIPs relative to input was analyzed using DESeq2^81^. RNAs with adjusted P values < 0.05 and fold enrichment > 1.5 were selected for Gene Ontology (GO) analysis.

GO biological process analysis was performed using the enrichGO function in clusterProfiler^82^ with human annotations from org.Hs.eg.db in R. All genes with valid adjusted P values in the DESeq2 results were supplied as the background. P values were adjusted using the Benjamini– Hochberg method^83^, with cutoffs of 0.1 for both unadjusted and adjusted P values and 0.2 for q values. Fold enrichment was calculated by dividing the proportion of genes assigned to each GO term in the RIP-enriched gene set by the corresponding proportion in the annotated background. GO terms were grouped into biological themes based on keyword matching of term descriptions. Results were plotted using ggplot2.

## Acknowledgements

We are grateful for technical assistance from Jadwiga Gasiorek and Eric Sounameto at University of Montreal, Christian Charbonneau at the bioimaging facility at University of Montreal, Annie Gosselin and Angelique Bellemare-Pelletier at Flow cytometry facility at University of Montreal, Gokul Varma at Northwestern University, and David Kirchenbuechler at the Microscopy facility at Northwestern. Primary human specimens were obtained from the Banque de Cellules Leucémiques du Québec (BCLQ), supported by the Cancer Research Network of the Fonds de Recherche du Québec - Santé (FRQS). We acknowledge support for these studies by the National Cancer Institute (R01 98571 and 80728 to KLBB and MLG), the content is solely the responsibility of the authors and does not necessarily represent the official views of the National Institutes of Health. These studies were also supported by the Leukemia and Lymphoma Society Canada and USA TRP R6513-20 (KLBB), Canada Research Chair in Molecular Biology of the Cell Nucleus to (KLBB), Hospira Foundation Professor of Translational Cancer Biology (KLBB), and Canadian Institutes for Health Research (PJT 159785, KLBB).

## Contributions

BCK, LMM, SP, NS designed, conducted experiments and analyzed data; AR, CE and EL analyzed data; WY, VTA, MCC, KA conducted experiments; SC and GJR provided reagents and reviewed the manuscript; MLG and KLBB designed experiments, analyzed data, wrote and edited the manuscript; BCK and LMM edited the manuscript.

## Supplemental Figure Legends

**Supplemental Figure 1. Characterization of AML cell lines with genetic manipulation of eIF4E. A.** Representative western blot from total cell lysates of MM6 and NOMO-1 cell lines demonstrating NOMO-1 cells have similar eIF4E levels to CD34+ cells from healthy donors, while MM6 cells have similar levels to high-eIF4E AML patient specimens. β-Actin is provided for loading control. Each lane refers to a different sample. **B**. Confocal micrograph of eIF4E nuclear and cytoplasmic staining in MM6 cells. Scale bar = 10 µm. **C, D.** Quantitation of western blots for NOMO-1 eIF4E relative to vector (Figure 1A) or MM6 CRISPR-4E relative to CRISPR-CTRL cells (Figure 1B, Supplemental Figure 1G). Each data point represents a biological replicate. Bar represents the mean. Standard deviation and p-values (Welch’s t test) are shown. **E.** Representative confocal micrograph in NOMO-1 eIF4E cells demonstrating that HA staining is specific as its signal is removed upon treatment with hyaluronidase (HAse). HA is red; DAPI is blue. **F.** Representative western blot of MM6 cells treated with the eIF4E inhibitor ribavirin or vehicle control demonstrating that ribavirin reduces eIF4E target proteins including Ezrin. β-Actin is provided as a loading control. **G, H**. Representative western blots demonstrating lower eIF4E levels and factors in the Ezrin-CD44-HA axis in MM6 and THP-1 CRISPR-4E cells compared to CRISPR-Controls.

**Supplemental Figure 2. Depletion of eIF4E reduces engraftment and improves overall survival in MM6 CDX model using an alternative CRISPR/Cas9 clone. A.** Schematic of the in vivo assay design. **B.** Validation that depletion of eIF4E decreases AML engraftment in bone marrow and spleen. Gating strategy for the evaluation of human AML cells (top panel). Percent engraftment in bone marrow (left bottom panel) and spleen (right bottom panel). Violin plot representing the median and quartiles, p-values calculated with a two tailed Mann-Whitney U test. **C.** Kaplan–Meier curves comparing overall survival of CDX mice with the indicated MM6 cell line clones. Log-rank (Mantel-Cox) test applied. **D.** Western blot analysis of AML cells isolated from leukemic mice demonstrating that CRISPR-4E MM6 cells have reduced eIF4E levels relative to CRISPR-CTRL, each lane represents a different mouse.

**Supplemental Figure 3. Depletion of eIF4E reduces leukemia engraftment in bone marrow and spleen using a THP-1 CDX model. A.** Schematic of the in vivo assay design. Engraftment in the bone marrow was assessed on day 30 and 59 post-transplantation. On day 56, a subset of mice (n = 3/cohort) was euthanized to evaluate leukemia engraftment in BM and spleen. Data for day 59 are presented in Figure 2. **B.** Engraftment analysis in BM aspirates form living mice at day 30 post-transplantation. Table shows the number of mice with detectable disease; quantification of THP-1 cells (frequencies) was not possible at this early stage due to their low levels. **C**. Engraftment analysis of a subset of mice at day 56 in bone marrow and spleen. Representative flow cytometry plots show AML cells in bone marrow and spleen from CRISPR-4E and CTRL groups (left panels). Percentage of human cells in the indicated organ is shown (right panels), each symbol represents an animal, bars represent the mean, standard deviation and p-values (Mann-Whitney U test) shown. **D.** Incidence of abdominal tumor formation at endpoint. Table shows the number of mice developing visible abdominal tumors in each group. **E.** Western blot analysis of AML cells isolated from leukemic mice confirming that THP-1 CRISPR-4E cells maintained reduced eIF4E levels compared with CRISPR-CTRL, each lane represents a different mouse.

**Supplemental Figure 4. Pharmacological inhibition of eIF4E impairs invasion and engraftment. A**. Invasion capacity of three high-eIF4E AML patient specimens relative to CD34+ cells from healthy donors. Each symbol reflects a different individual. Means and standard deviations, and fold change and p-value (two tailed Welch’s t test) per individual AML sample are shown. **B.** Representative flow cytometry plots illustrating how the frequency of AML cells was determined using staining with anti-human CD45 (hCD45) and anti-mouse CD45 (mCD45) antibodies. Graphs compare percentage of AML cells in peripheral blood in Ribavirin-treated mice (n = 5) and vehicle-treated controls (n = 5) at 111 and 118 days post-transplantation. Each symbol represents an animal, violin plot representing the median and quartiles, p-values calculated with a two tailed Mann-Whitney U test.

**Supplemental Figure 5. Cooperation between eIF4E and Ezrin. A.** Genetic reduction of Ezrin and eIF4E using siRNA in MM6 cells grown in suspension impacts production of factors in the Ezrin-CD44-HA axis compared to the siLUC control shown in a representative western blot (left panel). β-Actin is provided as loading control. Right panel, quantification of protein expression for the indicated siRNAs relative to siLUC is shown. The expression of each protein was calculated relative to β-Actin. Each symbol represents a biological replicate. Means, standard deviations and p-values (multiple paired t tests) are shown. **B.** Endogenous Ezrin and eIF4E immunoprecipitations (IP) in THP-1 total cell lysates show similar observations to MM6 cells (Figure 3C). SN supernatant after immunoprecipitation, IgG negative control. H2B also serves as a negative control for eIF4E and Ezrin IPs. **C.** Cytoplasmic (left) and total (right) cell lysates demonstrate Ezrin immunoprecipitated with the translation machinery including eIF4AI/II. **D.** Analysis of Ezrin IPs under native conditions and following RNase treatment, showing that RNase treatment did not affect the Ezrin–eIF4E interaction. **E.** Example fractionation controls for suspension (left) or invaded (right) cells shown in Figure 3F and G indicating quality of fractions. MEK and CalR are cytoplasmic markers; NOPP140 and H3K27A are nuclear markers. **F.** Count of the number of pseudopods per cell observed in suspension and invaded MM6 cells represented as a fraction relative to the total cells counted. Each symbol represents a biological replicate. Bars represent the mean, shown with standard deviations and p-values (two-way ANOVA).

**Supplemental Figure 6. Analysis of T-PODs. A.** Schema of VISTA-R for invaded cells. Steps for isolated invaded AML cells for marking active ribosomes with the VISTA-R method. **B.** Confocal microscopy for patient specimen AML23, demonstrated that active translation (characterized by VISTA-R) and Ezrin are present in the same pseudopods (white arrow). A single section through the plane of the cell is shown. Scale bar = 10µm. **C.** Representative western blot demonstrating that protein reduction due to siRNA to eIF4E or Ezrin occurs in invaded cells, thus invaded cells are not a result of rescue from the siRNA. Results are similar to MM6 cells grown in suspension (Figure 3A). **D.** Visualization of masks generated during Imaris confocal analysis to automate identification of pseudopods and subsequent measurement of contents. Scale bar = 10µm. **E.** Total rRNA levels detected by RT-qPCR in invaded MM6 cells as a function of genetic knockdown using siRNA to *EIF4E* or *EZR* as compared to siLUC treated cells. Each symbol represents a biological replicate performed independently. Bars represent the mean, shown with standard deviations and p-values (Welch’s t test). **F.** Representative western blots of invaded CRISPR MM6 cells showing reduced levels of factors in the Ezrin-CD44-HA axis similar to cells grown in suspension.

