## Supplemental Table 2 for "eIF4E and Ezrin drive a pseudopod-localized migratory translation program in AML"

**Name	Sequence**

**Mcl1Fw:** ACTTCTCACTTCCGCTTCCTTCCA

**Mcl1 Rv:** TTTGAGGCCAAACATTGCCAGTCG

**POL2AFw:** TGACTGCCAACACAGCCATCTACT

**POL2ARv:** GGGCCACATCAAAGTCAGGCATTT

**GAPDHFw:** GAAGGTGAAGGTCGGAGTC

**GAPDHRv:** GAAGATGGTGATGGGATTTC

**UbcFw:** att tgg gtc gcA gtt ctt g

**UbcRv:** tgc ctt gac attctc gat ggt

**ActinBF:** gcatggagtcctgtggcatccacg

**ActinBR:** ggtgtaacgcaactaagtcatag

**18S Fw:** CGG CGA CGA CCC ATT CGA AC

**18S Rv:** GAA TCG AAC CCT GAT TCC CCG TC

**RPL13aFw:** TTAATTCCTCATGCGTTGCCTGCC

**RPL13aRv:** TTCCTTGCTCCCAGCTTCCTATGT

**G6PDHF** TGGCAAAGTCGGTTTCTCTCTGGA

**G6PDHR** TTGGGAACATGTCTCAGACTGGCA

**HAS3Fw:** CAGGAGGACCCTGACTACTT

**HAS3Rv:** GTGGAAGATGTCCAGCATGTA

**EzrinFw:** CCGTGGGATGCTCAAAGATAA

**EzrinRv:** TCCAAGCCAAAGGTCTGTTC

**CD44Fw:** CGGCTCCTGTTAAATGGTATCT

**CD44Rv:** TCTGCTTTGTGGTCTGAGAAG

**UAP1Fw:** GCAGTGCTACAAGGGATCAA

**UAP1Rv:** CCACCAGCTAGAAGAAGAACTG

**eIF4EFw:** AGGAGGTTGCTAACCCAGAACACT

**eIF4ERv:** AAAGTGAGTAGTCACAGCCAGGCA

**HK2Fw:** TGTGAGGTCCACTCCAGAT

**Hk2Rv:** GAGCCCATTGTCCGTTACTT

**28SFw:** GTGCAGATCTTGGTGGTAGTAG

**28SRv:** TCCAAGCCAAAGGTCTGTTC

**5SFw:** GGCCATACCACCCTGAAC

**5SRv:** GGTCTCCCATCCAAGTACTAAC

**PGM5Fw:** TGATCTCCGAATCGACCTATCT

**PGM5Rv:** ATATCCACTGGGTCCACTATCT

**mtCox1Fw:** CAACCTCAACACCACCTTCT

**mtCox1Rv:** TCCGAAGCCTGGTAGGATAA

**mtCytBFw:** TCCCCACCCCATCCAACATCT

**mtCytBRv:** GGTAAAGAATCGTGTGAGGGT
