## Supplementary figures and images for "eIF4E and Ezrin drive a pseudopod-localized migratory translation program in AML"

### Supplemental Figure 1

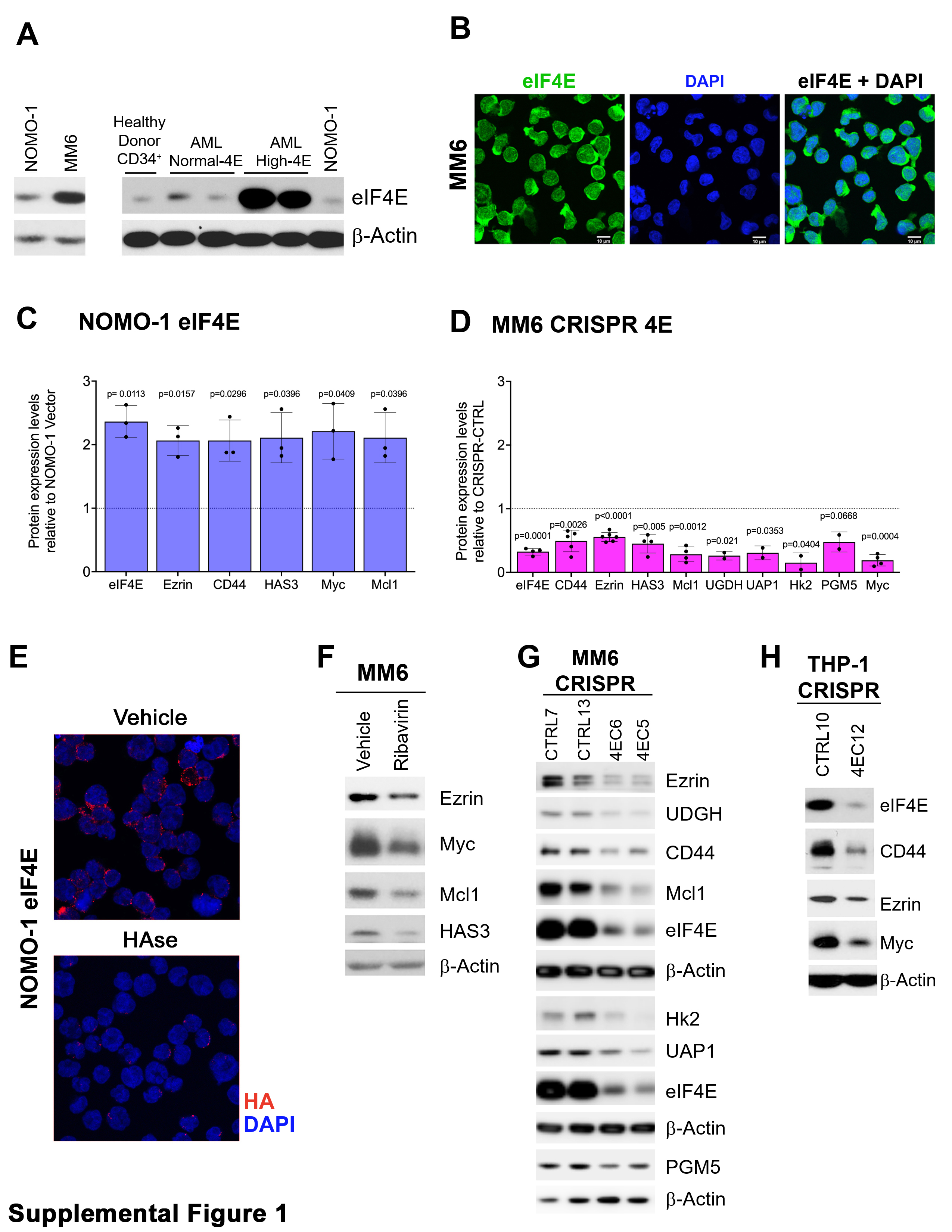

### Supplemental Figure 2

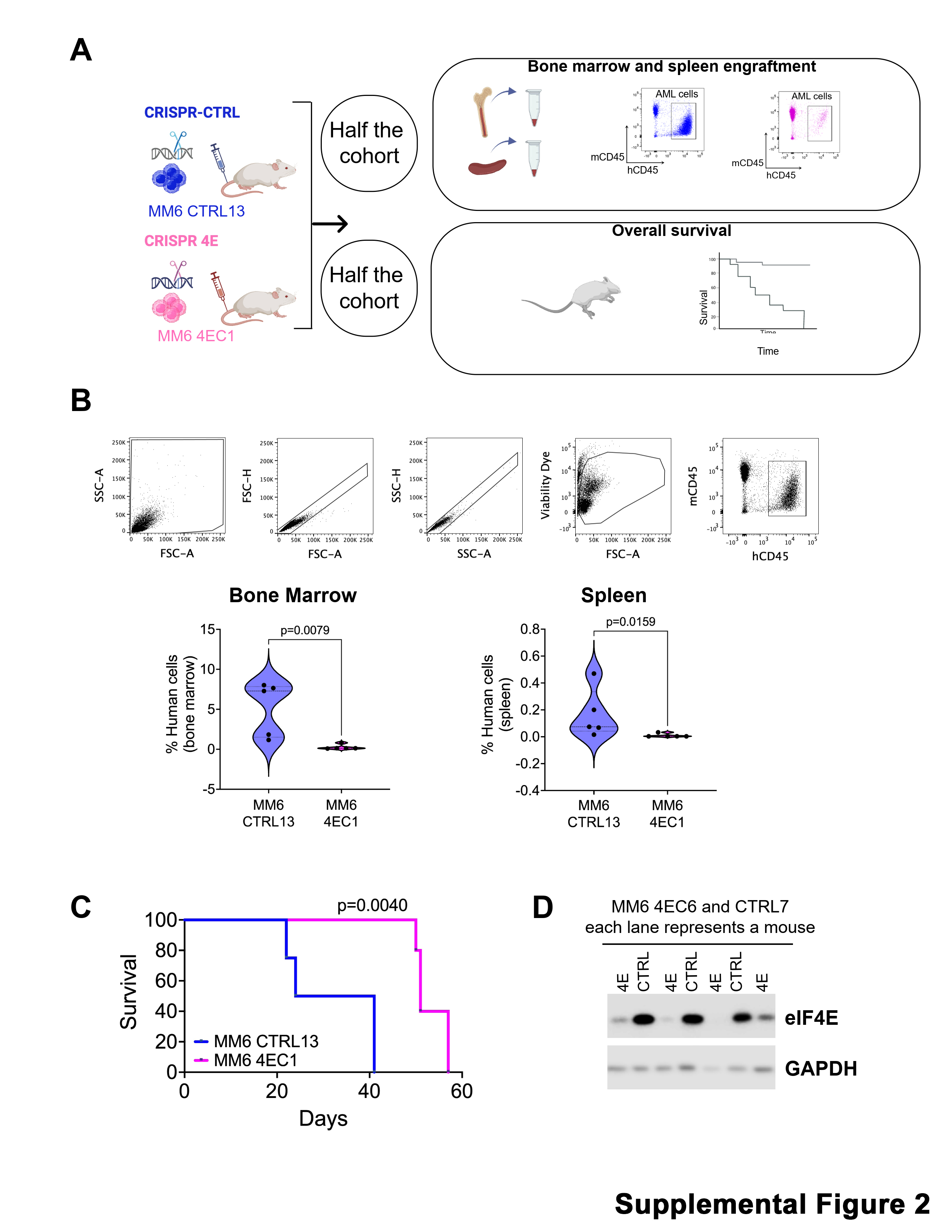

### Supplemental Figure 3

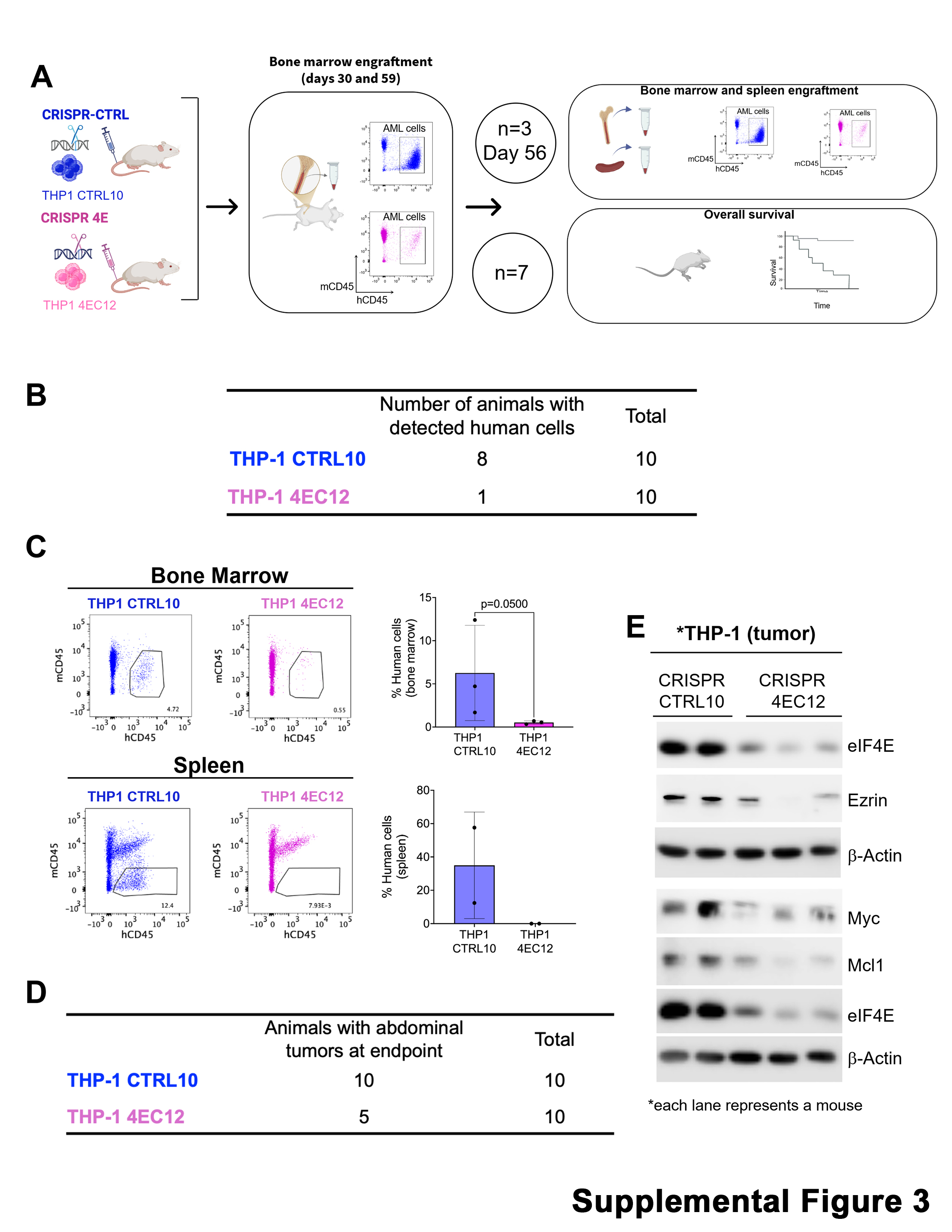

### Supplemental Figure 4

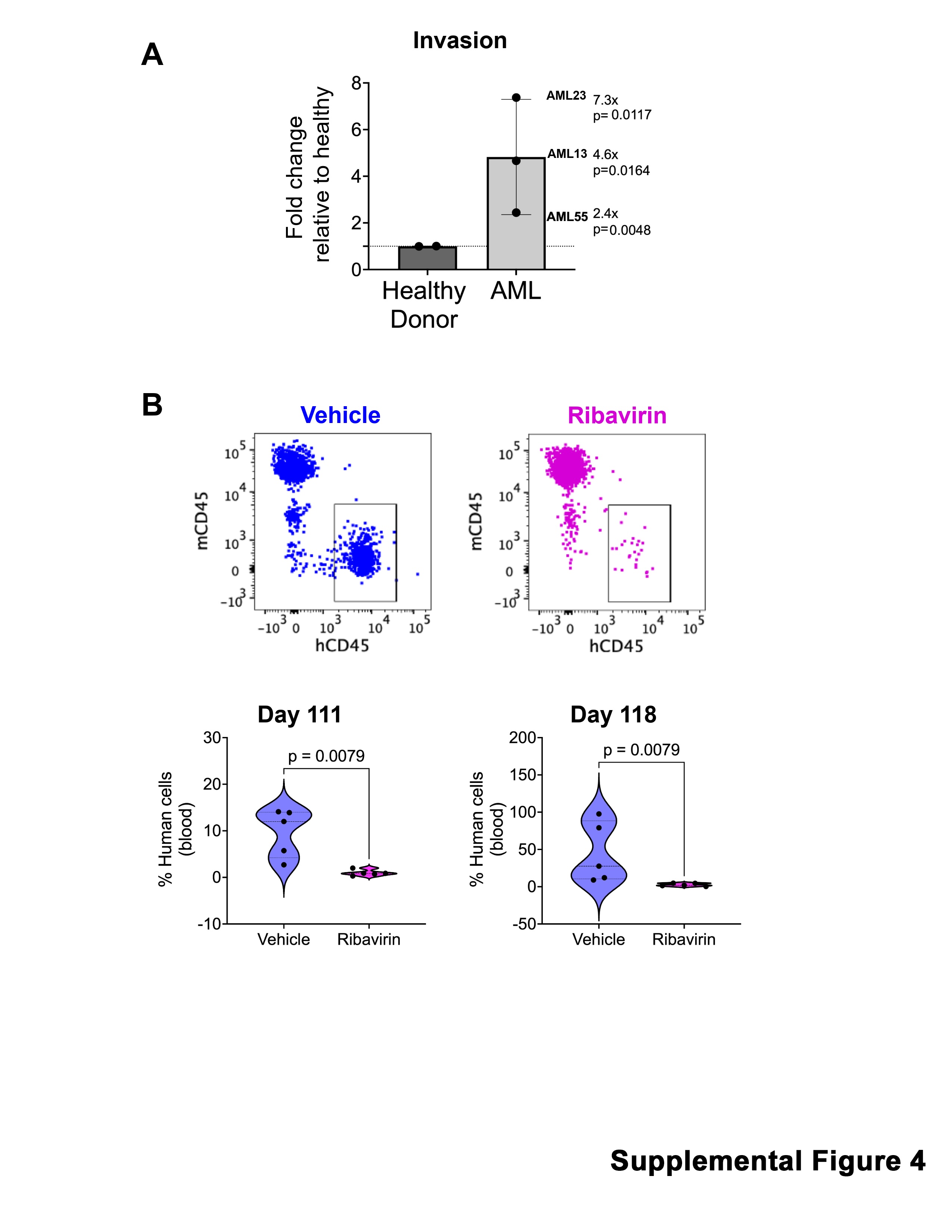

### Supplemental Figure 5

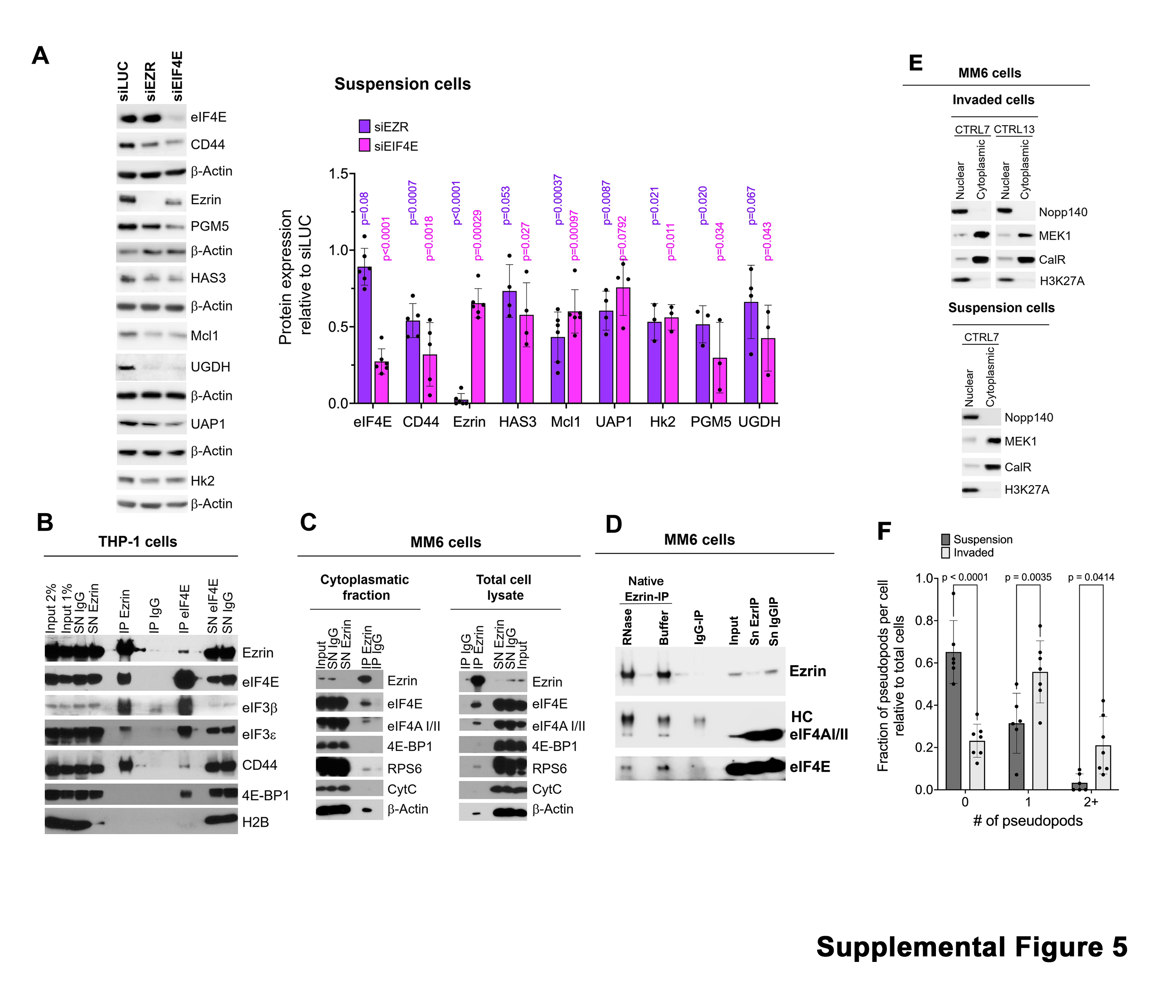

### Supplemental Figure 6

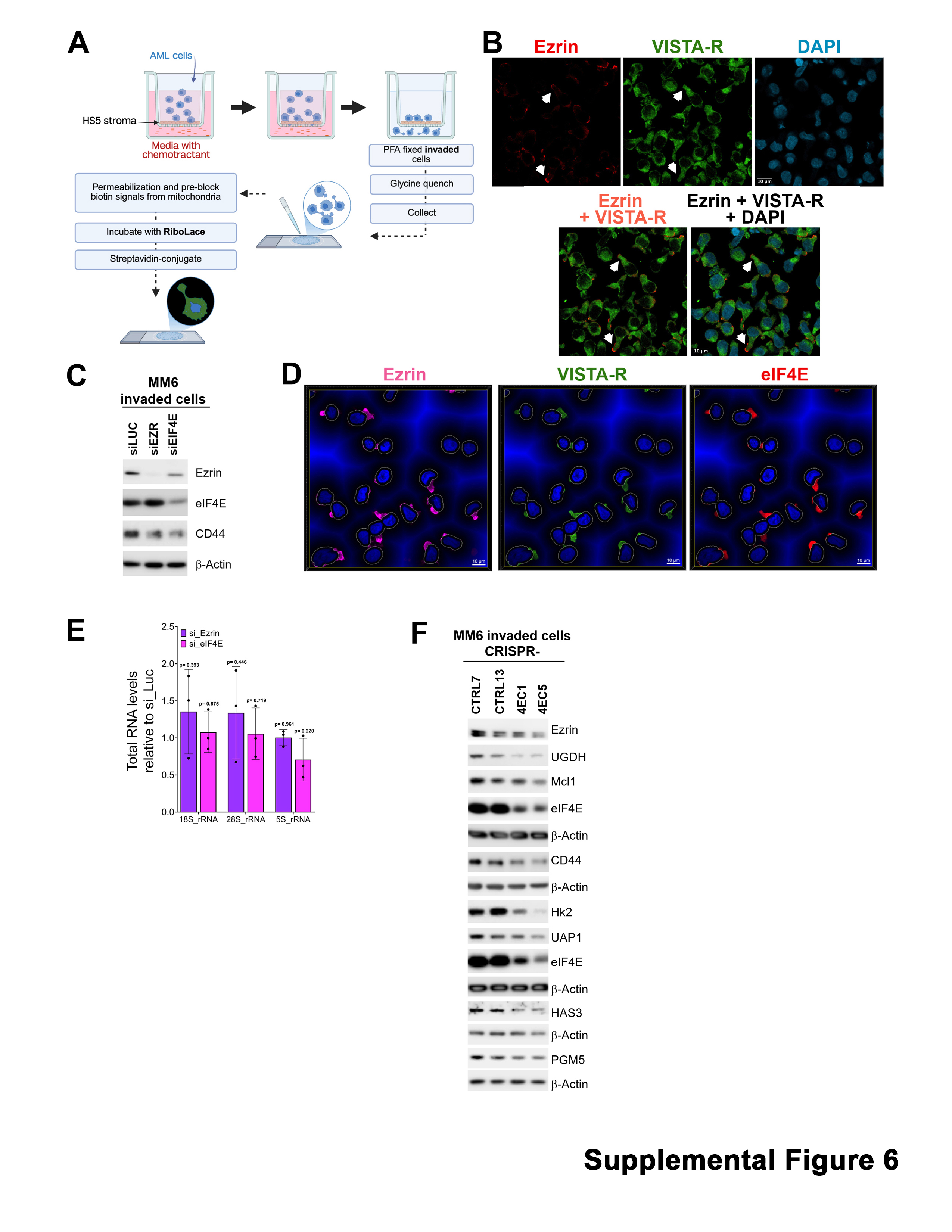
